# Normative modelling using deep autoencoders: a multi-cohort study on mild cognitive impairment and Alzheimer’s disease

**DOI:** 10.1101/2020.02.10.931824

**Authors:** Walter H. L. Pinaya, Cristina Scarpazza, Rafael Garcia-Dias, Sandra Vieira, Lea Baecker, Pedro F. da Costa, Alberto Redolfi, Giovanni B. Frisoni, Michela Pievani, Vince D. Calhoun, João R. Sato, Andrea Mechelli, the Alzheimer’s Disease Neuroimaging Initiative, the Australian Imaging Biomarkers and Lifestyle flagship study of ageing

**Author notes:** Corresponding author *Email address (Walter H. L. Pinaya). Data used in preparation of this article was obtained from the Alzheimer’s Disease Neuroimaging Initiative (ADNI) database (adni.loni.usc.edu). As such, the investigators within the ADNI contributed to the design and implementation of ADNI and/or provided data but did not participate in analysis or writing of this report. A complete listing of ADNI investigators can be found at: http://adni.loni.usc.edu/wp-content/uploads/how_to_apply/ADNI_Acknowledgement_List.pdf. Data used in the preparation of this article was obtained from the Australian Imaging Biomarkers and Lifestyle flagship study of ageing (AIBL) funded by the Commonwealth Scientific and Industrial Research Organisation (CSIRO) which was made available at the ADNI database (www.loni.usc.edu/ADNI).The AIBL researchers contributed data but did not participate in analysis or writing of this report. AIBL researchers are listed at www.aibl.csiro.au.

## Abstract

Normative modelling is an emerging method for quantifying how individuals deviate from the healthy populational pattern. Several machine learning models have been implemented to develop normative models to investigate brain disorders, including regression, support vector machines and Gaussian process models. With the advance of deep learning technology, the use of deep neural networks has also been proposed. In this study, we assessed normative models based on deep autoencoders using structural neuroimaging data from patients with Alzheimer’s disease (n=206) and mild cognitive impairment (n=354). We first trained the autoencoder on an independent dataset (UK Biobank dataset) with 11,034 healthy controls. Then, we estimated how each patient deviated from this norm and established which brain regions were associated to this deviation. Finally, we compared the performance of our normative model against traditional classifiers. As expected, we found that patients exhibited deviations according to the severity of their clinical condition. The model identified medial temporal regions, including the hippocampus, and the ventricular system as critical regions for the calculation of the deviation score. Overall, the normative model had comparable cross-cohort generalizability to traditional classifiers. In order to promote open science, we are making all scripts and the trained models available to the wider research community.

## 1. Introduction

Normative modelling is an emerging method for quantifying and describing how individuals deviate from the expected pattern learned from a population or large sample (Marquand et al., 2016). Recently, this approach has been applied to neuroimaging data to investigate a number of brain disorders, such as attention deficit hyperactivity disorder (Kia and Marquand, 2018; Wolfers et al., 2018), autism spectrum disorder (Pinaya et al., 2019; Zabihi et al., 2019), schizophrenia (Kia and Marquand, 2018; Pinaya et al., 2019; Wolfers et al., 2018) and dementia (Huizinga et al., 2018; Ziegler et al., 2014). The procedure of normative modelling used in these studies has two steps: (i) first, statistical models are estimated to characterise the typical brain data from a reference cohort; (ii) then, the estimated model is applied to a target clinical cohort in order to quantify the variation (e.g. due to the effect of brain disorders).

Many statistical models have been proposed for normative modelling, including regression, support vector machines and Gaussian process modelling (for an extensive list, see Marquand et al. 2019). In Pinaya et al. (2019), we proposed a normative modelling approach based on the use of deep autoencoders to evaluate psychiatric patients. The use of a deep learning approach (LeCun et al., 2015; Vieira et al., 2020) enables models to learn multiple levels of representation about the intricate structure of the data and identify the most important morphological characteristic of the healthy brain. In addition, in Pinaya et al. (2019), the models were able to detect deviations at the level of the individual, with patients with schizophrenia and patients with autism spectrum disorder presenting values significantly higher than the healthy controls (HC).

Similar to psychiatric disorders, the clinical interpretation of magnetic resonance imaging scans can be challenging in the context of neurodegenerative disorders, as brain alterations may be difficult to distinguish from those related to healthy ageing. The identification of disease-related alterations can be particularly tricky in the early stages of a disorder. For this reason, there is a grown interest in the development of methods for quantifying deviations of regional brain volumes that can discriminate between healthy and pathological ageing, with the ultimate aim of improving diagnostic and prognostic assessment of neurodegenerative disorders (Brewer, 2009). Here, we used the autoencoder normative method (Pinaya et al., 2019) to evaluate the most common type of dementia in the elderly worldwide, Alzheimer’s disease (AD).

First, we trained the normative models using a large number of HC subjects (*>*11,000 participants). Then, we assessed the performance of these models using data from patients with a diagnosis of mild cognitive impairment (MCI), the prodromal stage to AD, and patients with a diagnosis of AD. This assessment involved calculating the deviation, i.e. the extent to which subjects deviate from the norm, in five additional datasets composed of patients with MCI, patients with AD, and HC subjects. We had two main hypotheses. First, we hypothesised that the method would be robust and sensitive enough to create deviation values that reflect the severity of the brain anatomical alterations due to the disease, i.e. that individuals with AD would deviate from normality more than those with MCI. Second, we hypothesised that the main brain regions driving the observed deviation would include the medial temporal cortex and the ventricular system, consistent with the results of previous neuroimaging studies of MCI and AD (Busatto et al., 2008; Pini et al., 2016). Finally, we compared the performance of the normative approach against traditional classifiers to discriminate the patient groups from the HC group.

## 2. Methods

### 2.1. Datasets

In our analysis, we used six datasets: the UK Biobank (Sudlow et al., 2015), the Alzheimer’s Disease Neuroimaging Initiative (ADNI) (Mueller et al., 2005), the Australian Imaging Biomarkers and Lifestyle Study of Ageing (AIBL) (Ellis et al., 2009), the Alzheimer’s Disease Repository Without Borders (ARWiBo) (Frisoni et al., 2009; Galluzzi et al., 2010), the Open Access Series of Imaging Studies: Cross-Sectional (OASIS-1) (Marcus et al., 2007), and the Minimal Interval Resonance Imaging in Alzheimer’s Disease (MIRIAD) (Malone et al., 2013).

The UK Biobank is a study that aims to follow the health and well-being of 500,000 volunteer participants across the United Kingdom. From these participants, a subsample was chosen to collect multimodal imaging, including structural neuroimaging. Here, we used an early release of the project’s data comprising of 11,034 HC participants. The inclusion criteria for the present study were: a) subjects who had the data collected in the same MRI scanner (from Cheadle centre), b) age between 47 to 73 years old. The only exclusion criterion was previous hospitalization associated with the diagnosis of mental and behavioural disorders, disease of the nervous system, cerebrovascular diseases, benign neoplasm of meninges, brain and other parts of the central nervous system, or injuries to the head. More details about the dataset can be found elsewhere (Alfaro-Almagro et al., 2018; Elliott and Peakman, 2008; Miller et al., 2016; Sudlow et al., 2015).

The ADNI consortium started in 2003 as a public-private partnership, led by Principal Investigator Michael W. Weiner. Its goal was to verify whether different neuroimaging biomarkers and neuropsychological assessments can be combined to measure the progression of MCI and to study the development of AD (for more up-to-date information, see http://www.adni-info.org). In this study, we included the structural MRI collected during the ADNI GO, ADNI 2 and ADNI 3 phases. Similar to UK Biobank, we included only subjects with age between 47 to 73 years old. The final dataset comprised of 517 subjects, where 212 were HC, 159 were patients with early MCI (EMCI), 82 were patient with late MCI (LMCI), and 64 were patients with AD. In the ADNI datasets, participants were assigned to these MCI stages based on different levels of impairment on a single episodic memory measure, with the EMCI group showing milder episodic memory impairment than the LMCI group (Aisen et al., 2010; Edmonds et al., 2019).

The AIBL dataset was developed to enhance the understanding of the pathogenesis of AD, concentrating on its early diagnosis (more details can be found in Ellis et al. 2009). Here, we included the structural MRI of subjects between 47 to 73 years old, to match the age range of the UK Biobank dataset. The final group was composed of 346 subjects, where 262 were HC, 46 were patients with MCI (stage not known), and 38 were patients with AD.

The ARWiBo is a cross-sectional dataset including data from patients and controls enrolled at the Scientific Institute for the Research and Care of Alzheimer’s Disease [Istituto di Ricovero e Cura a Carattere Scientifico (IRCCS) Centro San Giovanni di Dio Fatebenefratelli, Brescia, Italy]. A multidisciplinary team of neurologists, neuroscientists, image analysists, neurophysiologists, and geneticists are involved in the assessment of patients. As part of their assessment, participants undergo blood drawing (for APOE genotyping), clinical and cognitive evaluations as well as high-resolution MRI scanning (more details can be found in Frisoni et al. 2009 and Galluzzi et al. 2010). Here, we included the structural MRI of subjects between 47 to 73 years old, to match the age range of the UK Biobank dataset. The resulting group was composed of 319 subjects, including 215 HC, 67 patients with MCI (stage not known), and 37 patients with AD.

The OASIS-1 dataset is the result of a collaborative effort of investigators from a single acquisition site supported by the National Institute on Aging (NIA), the Howard Hughes Medical Institute, the Biomedical Informatics Research Network (BIRN) and the Washington University Alzheimer’s Disease Research Center [Alzheimer’s Disease Research Center (ADRC)]. This collaborative effort aimed to create a freely available MRI dataset for the wider scientific community. The original dataset consisted of a cross-sectional collection of subjects aged 18 to 96. It included participants over the age of 60 who had received a clinical diagnosis of very mild to moderate AD (for more information, please see http://www.oasis-brains.org). In our analysis, we selected data collected from individuals who were between 47 to 73 years old, to match the age range of the UK Biobank dataset. The resulting group was composed of 78 subjects, including 41 HC and 37 patients with AD.

The MIRIAD dataset was designed to establish the minimal interval over which it would be feasible to undertake clinical trials in AD using atrophy measured from longitudinal MRI as an outcome measure (Malone et al., 2013). Here, we included the structural MRI of subjects between 47 to 73 years old, to match the age range of the UK Biobank dataset. The resulting group was composed of 48 subjects, including 18 HC and 30 patients with AD.

In the present study, we used the UK Biobank set to train the autoencoders and the ADNI, AIBL, ARWiBo, OASIS-1, and MIRIAD datasets to assess the normative model performance on data from patients with MCI and AD. To perform comparisons between HC and patient groups, we ensured that there were no significant statistical differences regarding age and sex in all three clinical datasets. We assessed each dataset independently using the ANOVA test to verify any differences in age and the Chi-square test of homogeneity to investigate differences in the sex ratios between groups (Table 1 and 2).

**Table 1:** Demographic information for the subjects from the UK Biobank dataset, the Alzheimer’s Disease Neuroimaging Initiative (ADNI) dataset, and the Australian Imaging Biomarkers and Lifestyle Study of Ageing (AIBL) dataset. We used ANOVA test and the chi-square test of homogeneity to test for significant differences in age and sex between healthy controls and patients. Abbreviations: HC = healthy control; EMCI = early mild cognitive impairment; LMCI = late mild cognitive impairment; AD = Alzheimer’s disease; MCI = mild cognitive impairment; SD = standard deviation.

|  | UK<br>BIOBANK<br>n=11,034 | ADNI<br>n=517 |  |  |  | p | AIBL<br>n=346 |  |  | p |
| --- | --- | --- | --- | --- | --- | --- | --- | --- | --- | --- |
|  |  | HC<br>n=212 | EMCI<br>n=159 | LMCI<br>n=82 | AD<br>n=64 |  | HC<br>n=262 | MCI<br>n=46 | AD<br>n=38 |  |
| Age, y |  |  |  |  |  | .87 |  |  |  | .28 |
| Mean±SD | 61.6±7.0 | 66.6±3.7 | 66.4±4.2 | 66.2±5.1 | 66.6±5.4 |  | 68.2±3.2 | 68.2±3.6 | 67.3±5.2 |  |
| Range | [47, 73] | [56, 72] | [56, 73] | [56, 73] | [56, 73] |  | [60, 73] | [56, 73] | [55, 73] |  |
| Sex, n (%) |  |  |  |  |  | .25 |  |  |  | .15 |
| Men | 5180 (47) | 90 (42) | 72 (45) | 40 (49) | 36 (56) |  | 113 (43) | 19 (41) | 17 (45) |  |
| Women | 5854 (53) | 122 (58) | 87 (55) | 42 (51) | 28 (44) |  | 149 (57) | 27 (59) | 21 (55) |  |

**Table 2:**
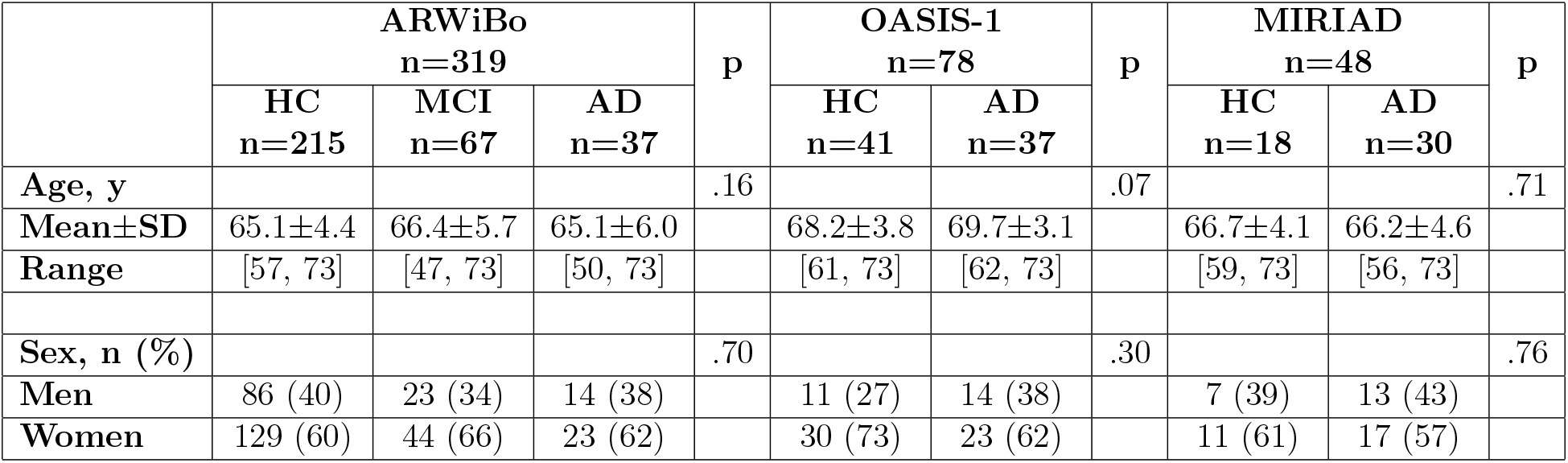
Demographic information for the subjects from the Alzheimer’s Disease Repository Without Borders (ARWiBo) dataset, the Open Access Series of Imaging Studies: Cross-Sectional (OASIS-1) dataset, and the Minimal Interval Resonance Imaging in Alzheimer’s Disease (MIRIAD) dataset. We used ANOVA test and the chi-square test of homogeneity to test for significant differences in age and sex between healthy controls and patients. Abbreviations: HC = healthy control; AD = Alzheimer’s disease; MCI = mild cognitive impairment; SD = standard deviation.

|  | ARWiBo<br>n=319 |  |  | p | OASIS-1<br>n=78 |  | p | MIRIAD<br>n=48 |  | p |
| --- | --- | --- | --- | --- | --- | --- | --- | --- | --- | --- |
|  | HC<br>n=215 | MCI<br>n=67 | AD<br>n=37 |  | HC<br>n=41 | AD<br>n=37 |  | HC<br>n=18 | AD<br>n=30 |  |
| **Age, y** |  |  |  | .16 |  |  | .07 |  |  | .71 |
| **Mean±SD** | 65.1±4.4 | 66.4±5.7 | 65.1±6.0 |  | 68.2±3.8 | 69.7±3.1 |  | 66.7±4.1 | 66.2±4.6 |  |
| **Range** | [57, 73] | [47, 73] | [50, 73] |  | [61, 73] | [62, 73] |  | [59, 73] | [56, 73] |  |
| **Sex, n (%)** |  |  |  | .70 |  |  | .30 |  |  | .76 |
| **Men** | 86 (40) | 23 (34) | 14 (38) |  | 11 (27) | 14 (38) |  | 7 (39) | 13 (43) |  |
| **Women** | 129 (60) | 44 (66) | 23 (62) |  | 30 (73) | 23 (62) |  | 11 (61) | 17 (57) |  |

### 2.2. MRI Processing

We used the FreeSurfer software (version 6.0) to estimate the brain regions’ volumes from the T1 weighted images. This estimation was performed using the “recon-all” command (see Fischl 2012 and Fischl et al. 2002, for more information). During this processing, the cortical surface of each hemisphere was parcellated according to the Desikan-Killiany atlas (Desikan et al., 2006) and anatomical volumetric measures were obtained via a wholebrain segmentation procedure (Aseg atlas) (Fischl et al., 2002). The final data included the cortical volume for each of the 68 cortical subregions (34 per hemisphere) and the volume of 33 neuroanatomical structures, totalling 101 subregions/structures (the complete list is presented in the supplementary materials).

### 2.3. Normative model

In this paper, we developed the normative model using the adversarial autoencoder (AAE; Figure 1) (Makhzani et al., 2015; Pinaya et al., 2020). As an autoencoder, this neural network has an encoder and a decoder. The function of the encoder is to take in an input *x* and map it into a latent encoding space, creating a latent code *h*. Then, the goal of the decoder is to reconstruct the input data based on the latent code. The AAE is a blend of this autoencoder framework with adversarial training, which is used in generative adversarial networks modelling (Goodfellow et al., 2014). This autoencoder uses the adversarial training to shape the distribution of the latent code to look similar to a predefined prior distribution. The AAE achieves this desired distribution by incorporating a discriminator network into its structure. In this scheme, the discriminator receives two types of inputs: random numbers sampled from the desired prior distribution, and the latent code. During the training process, the discriminator will make predictions regarding whether its input data was sampled from the prior distribution or the latent code. The adversarial training forces the encoder to produce a latent code space that can fool the discriminator into predicting that the encoded samples are just another sample from the prior distribution.

**Figure 1:**
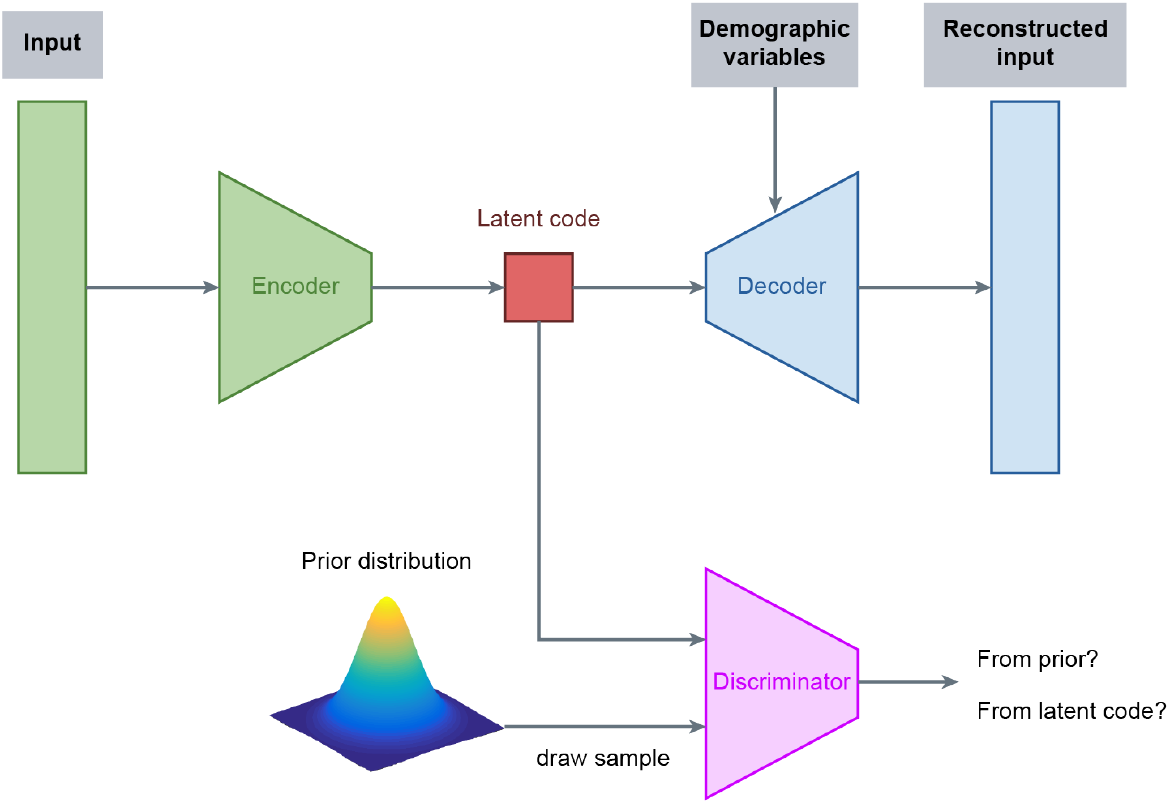
Structure of the normative model based on adversarial autoencoders. In this configuration, the subject data is inputted into the encoder and then mapped to the latent code. This latent code is fed to the decoder with the demographic data, and then the decoder generates a reconstruction of the original data. During the training of the model, the discriminator predicts if its input data came from the latent code or if it was randomly sampled from the chosen prior distribution (e.g. Gaussian distribution). Based on these predictions, the adversarial autoencoder forces the encoder to produce a latent code similar to the prior distribution selected. Since the model is trained on healthy controls data, it is expected that it can reconstruct similar data relatively well, yielding a small reconstruction error. However, the model is expected to generate a high error when processing data affected by unseen underlying mechanisms, e.g. pathological mechanisms.

In this study, we trained the AAE to codify and reconstruct the data of HC subjects. The main idea of this normative approach is that, since the AAE only learns how to reconstruct images from HC individuals, it will be less precise at mapping images from patients, which differ due to the pathological mechanisms of the disorder. As a result, the difference between the reconstructed data and the original data will be larger in patients than HC individuals. Regarding our model architecture, we used an encoder with two hidden layers with 100 neurons, and a latent code with a size of 20 neurons. The decoder and the discriminator had a similar structure (two hidden layers with 100 neurons). All hidden layers had a leaky ReLU non-linearity (Maas et al., 2013). The latent code and the decoder’s output layer had a linear activation function.

### 2.4. Normative model training

To train the autoencoder, first, we performed the pre-processing of the brain features. This involved estimating the relative brain region volumes for each subject by dividing the original brain region volumes by the total intracranial volume. Then, we normalised the relative brain region volumes across all the participants in the training set. In this step, we performed a normalisation robust to outliers by subtracting the median value of the relative brain region volume and then scaling the data according to its interquartile range. Centering and scaling was done independently for each brain region. The same relevant statistics (median and interquartile range) were later used to normalise the data from the clinical datasets before feeding them to the model.

In our analyses, we used a conditioned AAE (Makhzani et al., 2015). This type of autoencoder allows us to influence the model’s reconstruction using the demographic variables, i.e. age and sex. To input these variables into the model, we transformed age and sex into one-hot encoding vectors. After this transformation, each subject has an age vector with 27 positions, where each position corresponds to a year within the range of 47-73 years. In this vector, all positions have value zero except the one that indicates the subject’s age which has a value equal to 1. The subject’s sex was represented in a one-hot encoded vector with two positions, one for male and one for female. The AAE’s decoder used these vectors together with the latent code to reconstruct the brain data. This architecture forces the network to disentangle the label information from the latent code (Makhzani et al., 2015).

With the features pre-processed and the conditioning data prepared, we trained the autoencoder to minimise the mean squared value of its reconstruction error using Adam optimizer (Kingma and Ba, 2014) for 200 epochs. A minibatch approach was used in this gradient descent-based optimizer, with a batch size of 256. The model was trained with a cyclical learning rate (Smith, 2017), which allows convergence of the training with fewer epochs. We started using a base learning rate with a value of 0.0001 and a maximum learning rate value of 0.005, chosen using the “LR Range Test” (Smith, 2018). The learning rate cycle had a basic triangular shape with an amplitude decaying (gamma = 0.98).

In this study, we accessed the robustness of the autoencoder approach by training it with different simulated sets using the bootstrapping as the resampling method. We created 1,000 bootstrapped sets (each one with n = 11,032) by sampling with replacement from the UK Biobank. These bootstrapped sets were used to train the AAE. With this resampling method, we calculated: the value of the mean deviation (section 2.5) for each group from the ADNI, AIBL, ARWiBo, OASIS-1, and MIRIAD datasets, the discriminative performance of the normative approach (section 2.5), and the deviation from normality of each brain region (section 2.6).

### 2.5. Analysis of the observed deviations

Similar to Pinaya et al. (2019), we processed the data of each subject using the AAE, and we calculated the mean squared error between the reconstruction and the inputted data as the metric of brain deviation (Eq. 1).

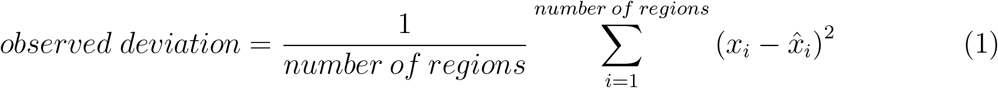

where *x*_*i*_ is the normalised value of the brain region *i*, 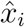 is the autoencoder reconstructed value of the brain region *i*, and number of regions is the number of cortical regions and neuroanatomical structures used (i.e. *number of regions* = 101).

In each iteration of the bootstrap method, we used the trained autoencoder to obtain the deviation metric of the subjects from the ADNI, AIBL, ARWiBo, OASIS-1, and MIRIAD datasets. Then, we calculated the difference between the mean deviation scores of each pair of groups. We identified a significant difference between groups if the confidence interval (95% of confidence) of this difference did not include the zero. Besides, we used the subjects’ deviations to obtain the discriminative performance of the autoencoder approach, measured by the area under the receiver operating characteristic curve (AUC).

### 2.6. Brain regions deviations

The autoencoder approach can quantify how much each brain region deviated from normality and contributed to the observed deviation. These values were obtained by measuring the difference between the inputted value and its reconstruction. In our study, we quantified the deviation for each subject from the ADNI, AIBL, ARWiBo, OASIS-1, and MIRIAD datasets. Then, in each iteration of the bootstrap method, we calculated the effect size of each brain region deviation – using Cliff’s delta (Cliff, 1993) value - between the HC group and each patient group.

### 2.7. Comparison against traditional machine learning classification

A further aim of the present study was to compare the performance of our normative model against a traditional classification approach. To measure the performance of the classifiers, we calculated the AUC using the .632+ bootstrap method (Efron and Tibshirani, 1997) with 1,000 iterations. Each clinical dataset (ADNI, AIBL, ARWiBo, OASIS-1, and MIRIAD) was analysed independently using the HC and patient groups to train the classifiers. Besides, the analysis was performed as multiple binary classifications between HC and each clinical group (e.g. HC versus LMCI).

In each iteration, first, we created the bootstrapped set by sampling the original data (from ADNI, AIBL, ARWiBo, OASIS-1, and MIRIAD datasets) with replacement. This bootstrapped set had the same size as the original dataset (for example, when analysing the ADNI dataset to classify healthy controls and patients with Alzheimer’s disease, the bootstrapped set had 212+64=276 subjects), and it contained repeated subjects (due to replacement). For each iteration, the subjects not included in the bootstrapped set were used as the out-of-bag set (i.e. test set).

Next, we obtained the relative brain region volumes of each subject by dividing the original volume by the total intracranial volume. Then, we normalised the values of the relative brain volumes across the subjects. In this normalisation step, we removed the median value of the brain regions and scaled the data according to the interquartile range. Centering and scaling was done independently for each brain region. The same relevant statistics (median and interquartile range) were later used to normalise the out-of-bag set.

To perform the classification analysis, we used a relevance vector machine (RVM) (Tipping, 2000) with a linear kernel. The RVM is a Bayesian treatment of identical functional form to the Support Vector Machines (SVM) (Cortes and Vapnik, 1995). One advantage of the RVM form over the SVM is that it is not necessary to estimate the error/margin trade-off parameter ’C’. After we trained the RVM on the bootstrapped set, we used the model to obtain the predicted probability of a subject belonging to the patient class. Using these probabilities, we calculated two AUC values, one for the bootstrapped set (called “resubstitution” metric) and one for the test set (called “out-of-bag” metric). By using the .632+ bootstrap method, we minimised the optimistic and pessimistic bias of the estimate and obtained the AUC value (Eq.2).

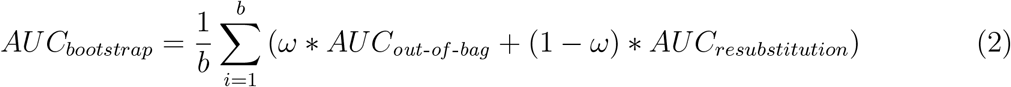

where *b* was the number of iterations and the weight *ω* was defined considering the relative overfitting rate (full description in Efron and Tibshirani 1997). To obtain the confidence interval (CI; 95% of confidence), we used the percentile method (Efron, 1981). Next, we compared these confidence intervals with the AUC obtained during the normative approach.

Finally, we compared the generalization of the classifiers with the results of the autoencoders. In this analysis, we used each trained classifier to predict the group of the subjects from the other clinical datasets. In order to verify if the performance in the independent datasets was significantly different from the normative approach, we calculated the difference between the AUCs of this generalization analysis and the AUCs of the autoencoders. With the 1,000 measures of the difference, we calculated its confidence interval (95% confidence) to verify if this difference is different from zero.

### 2.8. Experiments

We conducted our experiments in Python 3 using the Tensorflow 2.0 library (http://www.tensorflow.org/) and the sklearn rvm library (http://github.com/Mind-the-Pineapple/sklearn-rvm). We have made publicly available the codes and trained models used in this study at http://github.com/Warvito/Normative-modelling-using-deep-autoencoders. A Google’s Colaboratory notebook that calculates the deviations scores of new data is available at https://colab.research.google.com/github/Warvito/Normative-modelling-using-deep-autoencoders/blob/master/notebooks/predict.ipynb.

## 3. Results

### 3.1. Comparison of deviation values for healthy controls and patients

Figure 2 shows the mean value of the observed deviation for each group. For the ADNI dataset, we found a mean value of 0.28 ([0.27, 0.32]; 95% CI) for HC; 0.29 ([0.28, 0.35]; 95% CI) for EMCI; 0.32 ([0.30, 0.38]; 95% CI) for LMCI; 0.37 ([0.34, 0.47]; 95% CI) for AD. For the AIBL dataset, we found a mean value of 0.30 ([0.28, 0.33]; 95% CI) for HC; 0.36 ([0.34, 0.42]; 95% CI) for MCI; and 0.40 ([0.36, 0.50]; 95% CI) for AD. For the ARWiBo dataset, we found a mean value of 0.32 ([0.30, 0.38]; 95% CI) for HC; 0.37 ([0.34, 0.47]; 95% CI) for MCI; and 0.46 ([0.40, 0.62]; 95% CI) for AD. For the OASIS-1 dataset, we found a mean value of 0.41 ([0.39, 0.46]; 95% CI) for HC and 0.65 ([0.58, 0.79]; 95% CI) for AD. For the MIRIAD dataset, we found a mean value of 0.26 ([0.24, 0.30]; 95% CI) for HC and 0.48 ([0.41, 0.71]; 95% CI) for AD.

**Figure 2:**
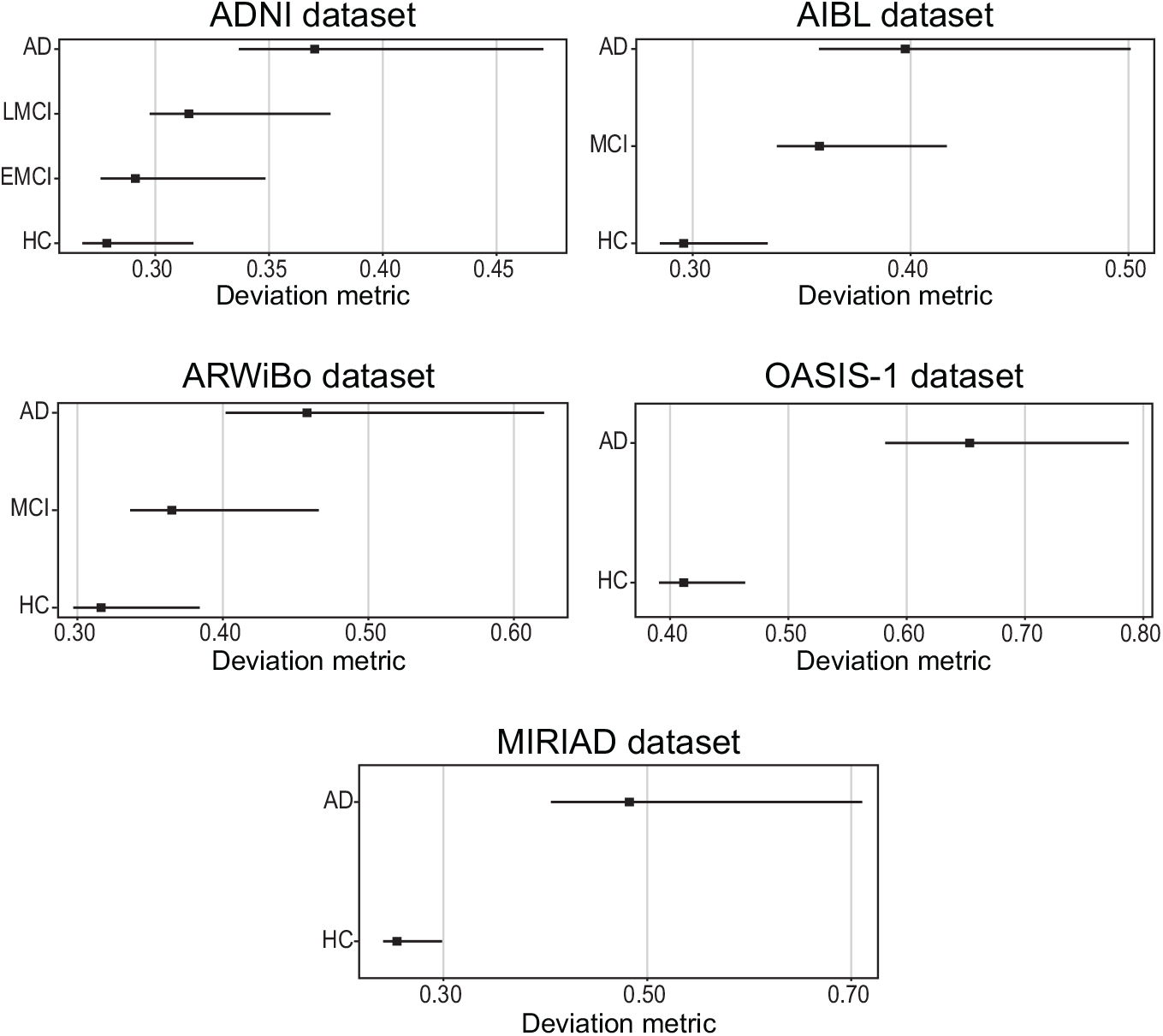
Mean value of the observed deviation calculated by the autoencoder for each group. The square marker indicates the mean value and the horizontal bars indicates the 95% confidence interval calculated using the percentile method on the bootstrap analysis. Abbreviations: AD = Alzheimer’s disease; EMCI = early mild cognitive impairment; LMCI = late mild cognitive impairment; MCI = mild cognitive impairment; HC = healthy controls; ADNI = Alzheimer’s Disease Neuroimaging Initiative; AIBL = Australian Imaging Biomarkers and Lifestyle Study of Ageing; ARWiBo = Alzheimer’s Disease Repository Without Borders; OASIS-1 = Open Access Series of Imaging Studies: Cross-Sectional; MIRIAD = Minimal Interval Resonance Imaging in Alzheimer’s Disease.

When we analysed the confidence interval of the difference between groups in the observed deviation, we obtained that, for the ADNI dataset, the difference between HC and EMCI was in the range [-0.03, 0.00], the difference between HC and LMCI was in the range [-0.06, −0.03], the difference between HC and AD group was in the interval of [-0.16, −0.06], the difference between EMCI and LMCI was in the range [-0.03, −0.02], the difference between EMCI and AD was in the range [-0.14, −0.06], and the difference between LMCI and AD was in the range [-0.10, −0.03]. For the AIBL dataset, the difference between HC and MCI was in the range [-0.09, −0.05], the difference between HC and AD was in the range [-0.17, −0.07], the difference between MCI and AD was in the range [-0.09, 0.00]. For the ARWiBo dataset, the difference between HC and MCI was in the range [-0.08, −0.03], the difference between HC and AD was in the range [-0.24, −0.10], the difference between MCI and AD was in the range [-0.16, −0.06]. For the OASIS-1 dataset, the difference between HC and AD was in the range [-0.18, −0.33]. Finally, for the MIRIAD dataset, the difference between HC and AD was in the range [-0.16, −0.41]. In summary, the five independent datasets presented mean deviation scores significantly different between their groups, except the comparison between HC and EMCI in the ADNI dataset and the comparison between MCI and AD in the AIBL dataset.

### 3.2. Normative model performance in discriminative tasks

We examined if the observed deviations were able to indicate if a person belonged to the patient or HC group (Figure 3). For the ADNI dataset, the normative approach had an AUC = 0.49 ([0.47, 0.51]; 95% CI) for patients with EMCI, an AUC = 0.61 ([0.58, 0.64]; 95% CI) for patients with LMCI, and an AUC = 0.74 ([0.70, 0.79]; 95% CI) for patients with AD. For the AIBL dataset, the normative approach had an AUC = 0.60 ([0.58, 0.64]; 95% CI) for patients with MCI, and an AUC = 0.71 ([0.65, 0.78]; 95% CI) for patients with AD. For the ARWiBo dataset, the normative approach had an AUC = 0.64 ([0.61, 0.68]; 95% CI) for patients with MCI, and an AUC = 0.84 ([0.78, 0.89]; 95% CI) for patients with AD. For the OASIS-1 dataset, the normative approach had an AUC = 0.72 ([0.67, 0.78]; 95% CI) for patients with AD. For the MIRIAD dataset, the normative approach had an AUC = 0.91 ([0.83, 0.97]; 95% CI) for patients with AD.

**Figure 3:**
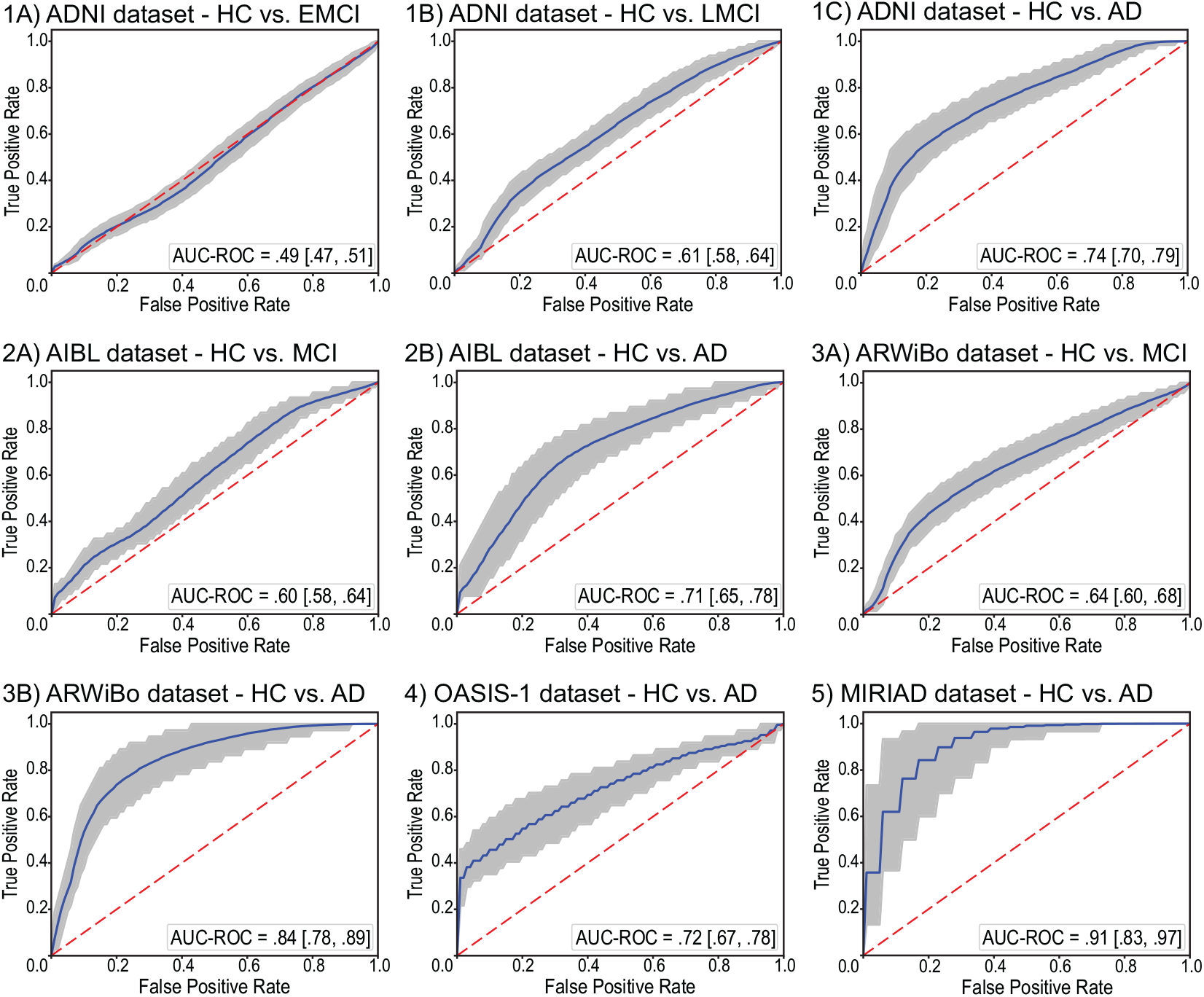
Discriminative performance of the normative approach. The solid line indicates the mean receiver operating characteristic curve across the bootstrap iterations with the shaded area indicating the 95% confidence interval calculated using the percentile method on the bootstrap analysis. The dashed line indicates the chance level. Abbreviations: AD = Alzheimer’s disease; AUC-ROC = area under the receiver operating characteristic curve; EMCI = early mild cognitive impairment; LMCI = late mild cognitive impairment; MCI = mild cognitive impairment; HC = healthy controls; ADNI = Alzheimer’s Disease Neuroimaging Initiative; AIBL = Australian Imaging Biomarkers and Lifestyle Study of Ageing; ARWiBo = Alzheimer’s Disease Repository Without Borders; OASIS-1 = Open Access Series of Imaging Studies: Cross-Sectional; MIRIAD = Minimal Interval Resonance Imaging in Alzheimer’s Disease.

### 3.3. Brain regions deviations

Figures 4 present the Cliff’s delta of each brain region when comparing its deviation in the HC group against the deviation in the patient groups. Only the regions with effect sizes significantly different from zero are shown (complete list presented in the supplementary materials). Among the regions showing significant deviation in patients with AD, we found the lateral ventricles, temporal horns, hippocampus, entorhinal cortex, parahippocampal cortex, and amygdala. A number of these regions also showed a high deviation in patients with MCI, including the lateral ventricles and hippocampus. Finally, we also noted that effect sizes were smaller for the regions identified in patients with MCI relative to those identified in patients with AD.

**Figure 4:**
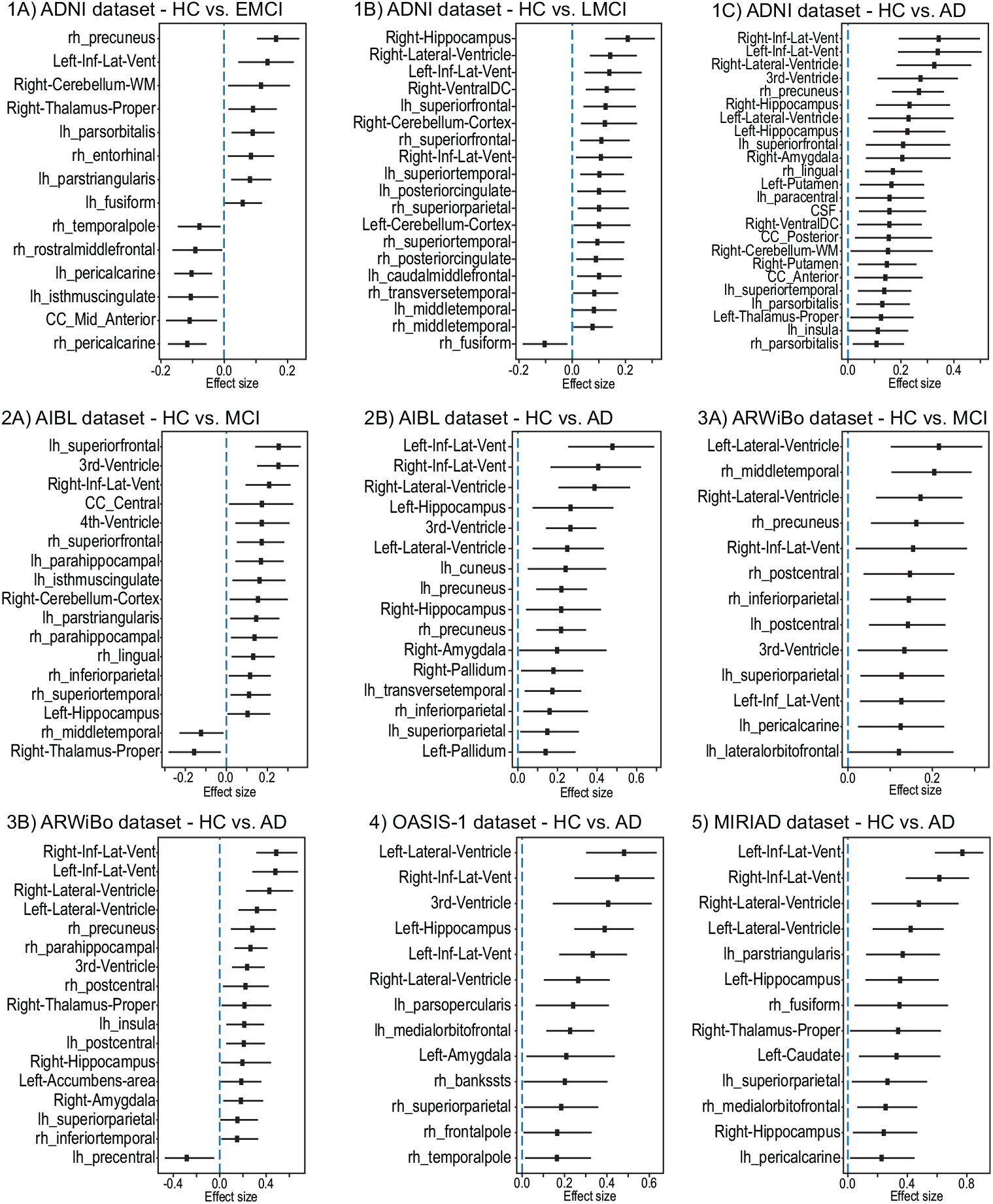
Brain regions deviations. The square marker indicates the mean effect size (Cliff’s delta) between the healthy control group and the respective patient groups. The horizontal bars indicate the 95% confidence interval calculated using the percentile method on the bootstrap analysis. Only the regions with a mean effect size significantly different from zero are presented. Abbreviations: AD = Alzheimer’s disease; AUCROC = area under the receiver operating characteristic curve; EMCI = early mild cognitive impairment; LMCI = late mild cognitive impairment; MCI = mild cognitive impairment; HC = healthy controls; ADNI = Alzheimer’s Disease Neuroimaging Initiative; AIBL = Australian Imaging Biomarkers and Lifestyle Study of Ageing; ARWiBo = Alzheimer’s Disease Repository Without Borders; OASIS-1 = Open Access Series of Imaging Studies: Cross-Sectional; MIRIAD = Minimal Interval Resonance Imaging in Alzheimer’s Disease.

### 3.4. Traditional machine learning classification

Using the RVM, we verified the performance of a traditional classifier when performing binary classification between HC and patients. For the ADNI dataset, we obtained an AUC = 0.69 ([0.58, 0.77]; 95% CI) when analysing patients with EMCI, an AUC = 0.76 ([0.64, 0.84]; 95% CI) when analysing patients with LMCI, and an AUC = 0.93 ([0.87, 0.97]; 95% CI) when analysing patients with AD. For the AIBL dataset, an AUC = 0.37 ([0.00, 0.78]; 95% CI) when analysing subjects with MCI, and we obtained an AUC = 0.93 ([0.86, 0.93]; 95% CI) when analysing patients with AD. Note, that the AUC for the AIBL dataset when analysing MCI had a wide interval. This interval was exacerbated due to the presence of overfitting and the .632+ bootstrap method compensatory effect that reduce the effect of bias caused by this overfitting. For the ARWiBo dataset, we obtained an AUC = 0.68 ([0.52, 0.78]; 95% CI) when analysing subjects with MCI, and an AUC = 0.94 ([0.87, 0.98]; 95% CI) when analysing patients with AD. For the OASIS-1 dataset, we obtained an AUC = 0.86 ([0.69, 0.96]; 95% CI) when analysing patients with AD. For the MIRIAD dataset, we obtained an AUC = 0.86 ([0.70, 0.96]; 95% CI) when analysing patients with AD.

To identify significant differences between the performance of the normative models and traditional classifiers, we calculated the confidence interval (95% of confidence) of the difference in AUC between the two methods. For the ADNI dataset we found that when classifying HC and EMCI the difference was in the range [-0.28, −0.09], when classifying HC and LMCI the difference was in the range [-0.24, −0.04], and when classifying HC and AD the difference was in the range [-0.25, −0.12]. For the AIBL dataset, when classifying the HC and MCI the difference of performance was in the range [-0.17, 0.74], and when classifying HC and AD the difference was in the range [-0.29, −0.12]. For the ARWiBo dataset, when classifying the HC and MCI the difference of performance was in the range [-0.15, 0.12], and when classifying HC and AD the difference was in the range [-0.17, 0.00]. For the OASIS-1 dataset, when classifying HC and AD the difference was in the range [-0.25, 0.04]. Finally, For the MIRIAD dataset, when classifying HC and AD the difference was in the range [-0.15, 0.06]. In summary, the traditional classifiers were superior to the normative models when predicting the difference between the groups in the ADNI dataset and the difference between HC and AD for the AIBL dataset; in contrast the performance of the two approaches was comparable for all other comparisons.

Finally, we examined how a classifier trained on a certain dataset would perform when applied to other datasets (i.e. cross-cohort generalizability). The results of this examination are presented in Table 3 and Table 4. When predicting AD, the classifiers had a higher mean performance than the normative approach in most cases (except when the model was trained on MIRIAD dataset and evaluated on ARWiBo dataset). However, the difference was not significantly different in almost half of the cases. When predicting MCI, the classifiers presented a lower mean performance in all cases, but the difference was not significantly different.

**Table 3:** Generalization performance of the classifiers for the classification between HC and patients with Alzheimer’s disease. In this table, the rows indicate the dataset where the classifier is trained and the columns indicate the dataset where the performance was tested. The area under the receiver operating characteristic curve is shown with the upper and lower bound of its 95% confidence interval. Performance significantly different from the normative approach calculated using the confidence interval of the difference between the approach across the bootstrap scheme is indicated by “*”.

|  | ADNI | AIBL | ARWiBo | OASIS-1 | MIRIAD |
| --- | --- | --- | --- | --- | --- |
| **ADNI** | - | 0.89 [0.93, 0.83]* | 0.88 [0.81, 0.93] | 0.84 [0.76, 0.90]* | 0.98 [0.95, 1.00] |
| **AIBL** | 0.88 [0.82, 0.93]* | - | 0.89 [0.93, 0.83]* | 0.86 [0.80, 0.91]* | 0.98 [0.94, 1.00] |
| **ARWiBo** | 0.88 [0.81, 0.92]* | 0.88 [0.83, 0.92]* | - | 0.80 [0.72, 0.86] | 0.96 [0.91, 0.99] |
| **OASIS-1** | 0.90 [0.84, 0.93]* | 0.89 [0.83, 0.93]* | 0.86 [0.77, 0.92] | - | 0.96 [0.91, 1.00] |
| **MIRIAD** | 0.89 [0.83, 0.93]* | 0.87 [0.80, 0.91]* | 0.82 [0.73, 0.91] | 0.83 [0.74, 0.89] | - |

**Table 4:** Generalization performance of the classifiers for the classification between HC and patients with mild cognitive impairment. In this table, the rows indicate the dataset where the classifier is trained and the columns indicate the dataset where the performance was measured. The area under the receiver operating characteristic curve is shown with the upper and lower bound of its 95% confidence interval. No case had a performance significantly different from the normative approach calculated using the confidence interval of the difference between the approach across the bootstrap scheme.

|  | AIBL | ARWiBo |
| --- | --- | --- |
| **AIBL** | - | 0.61 [0.54, 0.67] |
| **ARWiBo** | 0.59 [0.53, 0.65] | - |

## 4. Discussion

In this study, we evaluated the performance of the normative modelling approach based on deep autoencoders on data from patients with MCI and AD. Consistent with our first hypothesis, we found that the approach was effective in generating deviation values that reflect the severity of the disease, with patients with AD showing higher deviations than patients with MCI, and patients with LMCI showing larger deviations than patients with EMCI. We also measured how much each brain region deviated from normality and contributed to the observed deviation. Here, we found that regions from the ventricular system and medial temporal lobe were among those making the greatest significant contribution to deviation, consistent with our second hypothesis. Finally, we compared the performance of the normative approach versus a traditional classification approach. Although a higher performance was found for traditional classifiers in most cases, the difference was not statistically significant in the majority of cases.

We have replicated previous findings that the autoencoder is capable of detecting neuroanatomical deviation in individuals with brain disorders (Pinaya et al., 2019). In particular, in each of our five independent datasets, the normative model was able to assign higher values to patients with AD than healthy controls. This pattern was expected since the disorder is associated with profound alterations in the brain morphometry which were not present in the training set (Busatto et al., 2008; Pini et al., 2016). In addition, we have expanded these findings by demonstrating for the first time that autoencoders are capable of discriminating between different stages of the disease progress (i.e. EMCI versus LMCI versus AD). In particular, we observed that the MCI group presented intermediary deviation values in three independent datasets (ADNI, AIBL and ARWiBo). These values were also expected since the MCI is considered as a transitory stage between HC and AD (Morris et al., 2001), and usually present less brain atrophy compared to AD (Pihlajamaki et al., 2009). In addition, within the ADNI dataset, the MCI subjects were divided into two categories, EMCI and LMCI. Although individuals in both stages meet the conventional criteria for MCI, EMCI is associated with less pronounced symptoms thought to reflect an earlier point in the clinical spectrum than LMCI. In our analyses, we found that the patients with LMCI had a significantly (i.e. the confidence interval of the difference between the group do not overlap zero) larger deviation than patients with EMCI providing further confirmation that that deep autoencoders are capable of discriminating between different stages of the disease course.

With the autoencoder based approach, it was possible to identify the brain regions with the highest deviations from the expected normative values. Consistent with our second hypothesis, the AD group showed high levels of deviation in structures that are part of the ventricular system (such as the lateral ventricles, temporal horns, and 3rd ventricle) and in the medial temporal cortex, including the hippocampus, entorhinal cortex, parahippocampal cortex, and amygdala. Progressive ventricular expansion is one of the most reliable morphological changes in dementia patients, reflecting the increasing atrophy of the brain (Thompson et al., 2004). Likewise, medial temporal cortex atrophy is among the most consistent findings in neuroimaging studies of AD (Busatto et al., 2008; Fox and Schott, 2004) and an established marker of AD (Drago et al., 2011). While deviations in the MCI group had a smaller sizes than those in the AD group, there was a high degree of overlap in the hippocampus, parahippocampal cortex and several temporoparietal regions, consistent with previous neuroimaging studies of MCI (Chételat et al., 2002; Hämäläinen et al., 2007; Kang et al., 2019; Pennanen et al., 2005). The smaller effect size in MCI might be explained by two (not mutually-exclusive) factors: i) earlier stage in the AD course, hence milder atrophy, ii) heterogeneity of the MCI construct. Since MCI patients were not selected based on AD biomarkers (i.e., presence of beta-amyloid and tau protein in the cerebrospinal fluid) (Mulder et al., 2010), this group will likely include a mixture of AD and non-AD cases, hence the milder/diluted effect.

Finally, we compared the performance of our normative approach with traditional classifiers. The performance of the classifiers was measured in two schemes, on data from the same dataset where the model was trained and on data from independent clinical datasets (generalization performance). Although the traditional classifiers had a better mean performance in most cases, the differences between the two approaches were not statistically significant in most of the cases, especially when predicting the subjects from the ARWiBo, OASIS-1 and MIRIAD datasets. This similarity was more evident during the prediction of the patients with MCI (with exception the ADNI dataset).

Different from a case-control context, the normative approach does not need to be trained in a dataset with reasonable balancing between HC and patient groups. It is trained using only healthy controls, which enables the use of large cohorts of HC participants (Marquand et al., 2016, 2019), such as UK Biobank and Human Connectome Project (Van Essen et al., 2013). However, different from traditional classifiers, our normative approach cannot specify the diagnosis of the analysed subjects. Rather than seeing normative approaches as an alternative to classifiers, both methods could be used cooperatively to obtain computational systems with more reliable predictions. In order to promote open science, we have made all scripts and the trained models available to the wider research community (http://github.com/Warvito/Normative-modelling-using-deep-autoencoders).

## Acknowledgements

This study was supported by a Wellcome Trust’s Innovator Award to Andrea Mechelli (208519/Z/17/Z). This work was carried out within the scope of the project ”use-inspired basic research”, for which the Department of General Psychology of the University of Padova has been recognized as ”Dipartimento di eccellenza” by the Ministry of University and Research. JRS was supported by grant 2018/04654-9, São Paulo Research Foundation (FAPESP); and grant 2018/21934-5, São Paulo Research Foundation (FAPESP). VC was supported by NIH RF1AG063153.

This research has been conducted using the UK Biobank Resource (Project number: 40323).

Data were provided in part by OASIS: Cross-Sectional: Principal Investigators: D. Marcus, R, Buckner, J, Csernansky J. Morris; P50 AG05681, P01 AG03991, P01 AG026276, R01 AG021910, P20 MH071616, U24 RR021382.

Data collection and sharing for this project was funded by the Alzheimer’s Disease Neuroimaging Initiative (ADNI) (National Institutes of Health Grant U01 AG024904) and DOD ADNI (Department of Defense award number W81XWH-12-2-0012). ADNI is funded by the National Institute on Aging, the National Institute of Biomedical Imaging and Bioengineering, and through generous contributions from the following: AbbVie, Alzheimer’s Association; Alzheimer’s Drug Discovery Foundation; Araclon Biotech; BioClinica, Inc.; Biogen; Bristol-Myers Squibb Company; CereSpir, Inc.; Cogstate; Eisai Inc.; Elan Pharmaceuticals, Inc.; Eli Lilly and Company; EuroImmun; F. Hoffmann-La Roche Ltd and its affiliated company Genentech, Inc.; Fujirebio; GE Healthcare; IXICO Ltd.; Janssen Alzheimer Immunotherapy Research & Development, LLC.; Johnson & Johnson Pharmaceutical Research & Development LLC.; Lumosity; Lundbeck; Merck & Co., Inc.; Meso Scale Diagnostics, LLC.; NeuroRx Research; Neurotrack Technologies; Novartis Pharmaceuticals Corporation; Pfizer Inc.; Piramal Imaging; Servier; Takeda Pharmaceutical Company; and Transition Therapeutics. The Canadian Institutes of Health Research is providing funds to support ADNI clinical sites in Canada. Private sector contributions are facilitated by the Foundation for the National Institutes of Health (www.fnih.org). The grantee organization is the Northern California Institute for Research and Education, and the study is coordinated by the Alzheimer’s Therapeutic Research Institute at the University of Southern California. ADNI data are disseminated by the Laboratory for Neuro Imaging at the University of Southern California.

ARWiBo data (www.arwibo.it) was obtained from NeuGRID4You initiative funded by the European Commission (FP7/2007-2013) under grant agreement no.283562. The overall goal of ARWiBo is to contribute, thorough synergy with neuGRID (https://neugrid2.eu), to global data sharing and analysis in order to develop effective therapies, prevention methods and a cure for Alzheimer’ and other neurodegenerative diseases.

Data used in the preparation of this article were obtained from the MIRIAD database. The MIRIAD investigators did not participate in analysis or writing of this report. The MIRIAD dataset is made available through the support of the UK Alzheimer’s Society (Grant RF116). The original data collection was funded through an unrestricted educational grant from GlaxoSmithKline (Grant 6GKC).

## Author contributions

WHLP obtained and organized the MRI data, pre-processed the MRI images, implemented the normative models, performed the groups analysis, drafted and edited the manuscript; CS obtained and organized the MRI data and revised the manuscript; RGD implemented script to help pre-process the MRI data and the revised the manuscript; SV revised the manuscript; LB revised the manuscript; PFC implemented the traditional classifiers and revised the manuscript; AR was responsible for part of the data collection and revised the manuscript; MP was responsible for part of the data collection and revised the manuscript; GBF was responsible for part of the data collection; VDC revised manuscript and involved in funding efforts; JRS revised manuscript and involved in funding efforts; AM revised the manuscript, gathered funding, and supervised the project.

## Notes

https://github.com/Warvito/Normative-modelling-using-deep-autoencoders

