## supplementary materials for "Normative modelling using deep autoencoders: a multi-cohort study on mild cognitive impairment and Alzheimer’s disease"

[i] The Mind Research Network, NM, Albuquerque, USA.

[j] Tri-institutional Center for Translational Research in Neuroimaging and Data Science (TReNDS) [Georgia State, Georgia Tech, Emory], Atlanta, Georgia, USA.

** Data used in preparation of this article was obtained from the Alzheimer's Disease Neuroimaging Initiative (ADNI) database (adni.loni.usc.edu). As such, the investigators within the ADNI contributed to the design and implementation of ADNI and/or provided data but did not participate in analysis or writing of this report. A complete listing of ADNI investigators can be found at: http://adni.loni.usc.edu/wp-content/uploads/how_to_apply/ADNI_Acknowledgement_List.pdf

† Data used in the preparation of this article was obtained from the Australian Imaging Biomarkers and Lifestyle flagship study of ageing (AIBL) funded by the Commonwealth Scientific and Industrial Research Organisation (CSIRO) which was made available at the ADNI database (www.loni.usc.edu/ADNI). The AIBL researchers contributed data but did not participate in analysis or writing of this report. AIBL researchers are listed at www.aibl.csiro.au.

### 1. List of brain features

| | |
|---|---|
| Left-Lateral-Ventricle | lh_parstriangularis_volume |
| Left-Inf-Lat-Vent | lh_pericalcarine_volume |
| Left-Cerebellum-White-Matter | lh_postcentral_volume |
| Left-Cerebellum-Cortex | lh_posteriorcingulate_volume |
| Left-Thalamus-Proper | lh_precentral_volume |
| Left-Caudate | lh_precuneus_volume |
| Left-Putamen | lh_rostralanteriorcingulate_volume |
| Left-Pallidum | lh_rostralmiddlefrontal_volume |
| 3rd-Ventricle | lh_superiorfrontal_volume |
| 4th-Ventricle | lh_superiorparietal_volume |
| Brain-Stem | lh_superiortemporal_volume |
| Left-Hippocampus | lh_supramarginal_volume |
| Left-Amygdala | lh_frontalpole_volume |
| CSF | lh_temporalpole_volume |
| Left-Accumbens-area | lh_transversetemporal_volume |
| Left-VentralDC | lh_insula_volume |
| Right-Lateral-Ventricle | rh_bankssts_volume |
| Right-Inf-Lat-Vent | rh_caudalanteriorcingulate_volume |
| Right-Cerebellum-White-Matter | rh_caudalmiddlefrontal_volume |
| Right-Cerebellum-Cortex | rh_cuneus_volume |
| Right-Thalamus-Proper | rh_entorhinal_volume |
| Right-Caudate | rh_fusiform_volume |
| Right-Putamen | rh_inferiorparietal_volume |
| Right-Pallidum | rh_inferiortemporal_volume |
| Right-Hippocampus | rh_isthmuscingulate_volume |
| Right-Amygdala | rh_lateraloccipital_volume |
| Right-Accumbens-area | rh_lateralorbitofrontal_volume |
| Right-VentralDC | rh_lingual_volume |
| CC_Posterior | rh_medialorbitofrontal_volume |
| CC_Mid_Posterior | rh_middletemporal_volume |
| CC_Central | rh_parahippocampal_volume |
| CC_Mid_Anterior | rh_paracentral_volume |
| CC_Anterior | rh_parsopercularis_volume |
| lh_bankssts_volume | rh_parsorbitalis_volume |
| lh_caudalanteriorcingulate_volume | rh_parstriangularis_volume |
| lh_caudalmiddlefrontal_volume | rh_pericalcarine_volume |
| lh_cuneus_volume | rh_postcentral_volume |
| lh_entorhinal_volume | rh_posteriorcingulate_volume |
| lh_fusiform_volume | rh_precentral_volume |
| lh_inferiorparietal_volume | rh_precuneus_volume |
| lh_inferiortemporal_volume | rh_rostralanteriorcingulate_volume |
| lh_isthmuscingulate_volume | rh_rostralmiddlefrontal_volume |
| lh_lateraloccipital_volume | rh_superiorfrontal_volume |
| lh_lateralorbitofrontal_volume | rh_superiorparietal_volume |
| lh_lingual_volume | rh_superiortemporal_volume |
| lh_medialorbitofrontal_volume | rh_supramarginal_volume |
| lh_middletemporal_volume | rh_frontalpole_volume |
| lh_parahippocampal_volume | rh_temporalpole_volume |
| lh_paracentral_volume | rh_transversetemporal_volume |
| lh_parsopercularis_volume | rh_insula_volume |
| lh_parsorbitalis_volume | |

#### 2. Univariate analysis – ADNI dataset– HC vs EMCI

Table 1 - Statistical significance measured by the Mann-Whitney U test and effect size measured by Cliff's delta absolute value based on the comparison of the reconstruction error for each brain region between the HC and the EMCI groups from the ADNI dataset. The regions with p-value <= 0.05 are highlighted in bold.

| Regions | Effect size | p-value | Regions | Effect size | p-value |
|---|---|---|---|---|---|
| Left-Lateral-Ventricle | -0.045 | 0.230 | **lh_parstriangularis_volume** | **0.130** | **0.016** |
| **Left-Inf-Lat-Vent** | **-0.112** | **0.033** | lh_pericalcarine_volume | 0.046 | 0.226 |
| Left-Cerebellum-White-Matter | -0.039 | 0.260 | lh_postcentral_volume | 0.077 | 0.101 |
| Left-Cerebellum-Cortex | 0.094 | 0.060 | **lh_posteriorcingulate_volume** | **0.140** | **0.010** |
| **Left-Thalamus-Proper** | **0.209** | **>0.001** | lh_precentral_volume | 0.060 | 0.160 |
| Left-Caudate | 0.062 | 0.153 | **lh_precuneus_volume** | **0.085** | **0.081** |
| Left-Putamen | 0.025 | 0.339 | lh_rostralanteriorcingulate_volume | 0.095 | 0.059 |
| Left-Pallidum | -0.019 | 0.380 | **lh_rostralmiddlefrontal_volume** | **0.124** | **0.021** |
| 3rd-Ventricle | -0.021 | 0.365 | **lh_superiorfrontal_volume** | **0.148** | **0.007** |
| **4th-Ventricle** | **0.153** | **0.006** | lh_superiorparietal_volume | 0.047 | 0.220 |
| Brain-Stem | 0.090 | 0.069 | **lh_superiortemporal_volume** | **0.139** | **0.011** |
| Left-Hippocampus | 0.085 | 0.082 | lh_supramarginal_volume | 0.034 | 0.289 |
| Left-Amygdala | 0.079 | 0.097 | lh_frontalpole_volume | -0.071 | 0.121 |
| **CSF** | **-0.108** | **0.038** | **lh_temporalpole_volume** | **0.143** | **0.009** |
| Left-Accumbens-area | 0.008 | 0.447 | lh_transversetemporal_volume | 0.033 | 0.295 |
| Left-VentralDC | 0.038 | 0.265 | **lh_insula_volume** | **0.189** | **0.001** |
| Right-Lateral-Ventricle | -0.050 | 0.205 | rh_bankssts_volume | 0.005 | 0.466 |
| **Right-Inf-Lat-Vent** | **-0.103** | **0.046** | rh_caudalanteriorcingulate_volume | 0.006 | 0.460 |
| Right-Cerebellum-White-Matter | -0.065 | 0.142 | **rh_caudalmiddlefrontal_volume** | **0.132** | **0.015** |
| **Right-Cerebellum-Cortex** | **0.112** | **0.033** | rh_cuneus_volume | -0.031 | 0.306 |
| **Right-Thalamus-Proper** | **0.199** | **0.001** | rh_entorhinal_volume | 0.010 | 0.437 |
| Right-Caudate | 0.072 | 0.116 | rh_fusiform_volume | 0.062 | 0.153 |
| Right-Putamen | 0.010 | 0.437 | rh_inferiorparietal_volume | 0.065 | 0.141 |
| Right-Pallidum | -0.016 | 0.398 | rh_inferiortemporal_volume | 0.024 | 0.345 |
| **Right-Hippocampus** | **0.123** | **0.022** | rh_isthmuscingulate_volume | 0.068 | 0.130 |
| Right-Amygdala | 0.008 | 0.448 | rh_lateraloccipital_volume | -0.012 | 0.420 |
| **Right-Accumbens-area** | **-0.107** | **0.039** | **rh_lateralorbitofrontal_volume** | **0.091** | **0.067** |
| Right-VentralDC | 0.027 | 0.328 | rh_lingual_volume | -0.050 | 0.207 |
| CC_Posterior | -0.035 | 0.284 | **rh_medialorbitofrontal_volume** | **0.156** | **0.005** |
| CC_Mid_Posterior | 0.029 | 0.318 | **rh_middletemporal_volume** | **0.100** | **0.050** |
| **CC_Central** | **0.103** | **0.045** | rh_parahippocampal_volume | 0.029 | 0.315 |
| CC_Mid_Anterior | 0.031 | 0.304 | **rh_paracentral_volume** | **0.113** | **0.032** |
| CC_Anterior | -0.033 | 0.292 | **rh_parsopercularis_volume** | **0.109** | **0.036** |
| **lh_bankssts_volume** | **0.118** | **0.026** | rh_parsorbitalis_volume | 0.025 | 0.338 |
| lh_caudalanteriorcingulate_volume | 0.065 | 0.140 | rh_parstriangularis_volume | 0.075 | 0.109 |
| **lh_caudalmiddlefrontal_volume** | 0.031 | 0.306 | rh_pericalcarine_volume | 0.078 | 0.099 |
| **lh_cuneus_volume** | -0.041 | 0.248 | **rh_postcentral_volume** | **0.101** | **0.048** |
| **lh_entorhinal_volume** | **-0.092** | **0.065** | rh_posteriorcingulate_volume | 0.056 | 0.179 |
| lh_fusiform_volume | 0.052 | 0.195 | **rh_precentral_volume** | **0.105** | **0.041** |
| **lh_inferiorparietal_volume** | **0.100** | **0.049** | rh_precuneus_volume | 0.042 | 0.242 |
| **lh_inferiortemporal_volume** | **0.092** | **0.064** | **rh_rostralanteriorcingulate_volume** | **0.112** | **0.033** |
| lh_isthmuscingulate_volume | -0.048 | 0.213 | rh_rostralmiddlefrontal_volume | 0.084 | 0.083 |
| lh_lateraloccipital_volume | -0.002 | 0.484 | **rh_superiorfrontal_volume** | **0.114** | **0.030** |
| lh_lateralorbitofrontal_volume | 0.077 | 0.101 | rh_superiorparietal_volume | 0.030 | 0.308 |
| lh_lingual_volume | 0.007 | 0.457 | **rh_superiortemporal_volume** | **0.221** | **0.000** |
| **lh_medialorbitofrontal_volume** | **0.232** | **0.000** | rh_supramarginal_volume | 0.083 | 0.084 |
| **lh_middletemporal_volume** | **0.109** | **0.037** | **rh_frontalpole_volume** | **-0.104** | **0.043** |
| lh_parahippocampal_volume | -0.014 | 0.406 | **rh_temporalpole_volume** | **0.116** | **0.028** |
| **lh_paracentral_volume** | **0.126** | **0.019** | **rh_transversetemporal_volume** | **0.139** | **0.011** |
| **lh_parsopercularis_volume** | **0.134** | **0.014** | **rh_insula_volume** | **0.172** | **0.002** |
| lh_parsorbitalis_volume | -0.012 | 0.424 | | | |

#### 3. Univariate analysis – ADNI dataset – HC vs LMCI

Table 2 - Statistical significance measured by the Mann-Whitney U test and effect size measured by Cliff's delta absolute value based on the comparison of the reconstruction error for each brain region between the HC and the LMCI groups from the ADNI dataset. The regions with p-value <= 0.05 are highlighted in bold.

| Regions | Effect size | p-value | Regions | Effect size | p-value |
|---|---|---|---|---|---|
| **Left-Lateral-Ventricle** | **-0.106** | **0.080** | **lh_parstriangularis_volume** | **0.232** | **0.001** |
| **Left-Inf-Lat-Vent** | **-0.299** | **>0.001** | lh_pericalcarine_volume | 0.065 | 0.193 |
| Left-Cerebellum-White-Matter | -0.054 | 0.238 | **lh_postcentral_volume** | **0.191** | **0.006** |
| **Left-Cerebellum-Cortex** | **0.127** | **0.045** | **lh_posteriorcingulate_volume** | **0.132** | **0.040** |
| **Left-Thalamus-Proper** | **0.240** | **0.001** | **lh_precentral_volume** | **0.147** | **0.026** |
| Left-Caudate | 0.098 | 0.096 | **lh_precuneus_volume** | **0.246** | **0.001** |
| Left-Putamen | 0.098 | 0.096 | lh_rostralanteriorcingulate_volume | 0.113 | 0.066 |
| Left-Pallidum | 0.016 | 0.415 | **lh_rostralmiddlefrontal_volume** | **0.239** | **0.001** |
| **3rd-Ventricle** | **-0.161** | **0.016** | **lh_superiorfrontal_volume** | **0.287** | **>0.001** |
| 4th-Ventricle | 0.039 | 0.301 | lh_superiorparietal_volume | 0.120 | 0.055 |
| Brain-Stem | 0.068 | 0.183 | **lh_superiortemporal_volume** | **0.298** | **>0.001** |
| **Left-Hippocampus** | **0.366** | **>0.001** | **lh_supramarginal_volume** | **0.231** | **0.001** |
| **Left-Amygdala** | **0.278** | **>0.001** | lh_frontalpole_volume | 0.123 | 0.051 |
| **CSF** | **-0.245** | **0.001** | **lh_temporalpole_volume** | **0.142** | **0.029** |
| **Left-Accumbens-area** | **0.222** | **0.002** | lh_transversetemporal_volume | 0.086 | 0.126 |
| Left-VentralDC | 0.112 | 0.068 | **lh_insula_volume** | **0.254** | **>0.001** |
| Right-Lateral-Ventricle | -0.106 | 0.079 | **rh_bankssts_volume** | **0.227** | **0.001** |
| **Right-Inf-Lat-Vent** | **-0.258** | **>0.001** | rh_caudalanteriorcingulate_volume | 0.001 | 0.493 |
| Right-Cerebellum-White-Matter | -0.108 | 0.076 | **rh_caudalmiddlefrontal_volume** | **0.142** | **0.029** |
| Right-Cerebellum-Cortex | 0.118 | 0.058 | rh_cuneus_volume | 0.079 | 0.147 |
| **Right-Thalamus-Proper** | **0.336** | **>0.001** | **rh_entorhinal_volume** | **0.214** | **0.002** |
| Right-Caudate | 0.066 | 0.192 | **rh_fusiform_volume** | **0.198** | **0.004** |
| Right-Putamen | 0.087 | 0.125 | **rh_inferiorparietal_volume** | **0.345** | **>0.001** |
| Right-Pallidum | -0.044 | 0.280 | **rh_inferiortemporal_volume** | **0.267** | **>0.001** |
| **Right-Hippocampus** | **0.382** | **>0.001** | rh_isthmuscingulate_volume | 0.106 | 0.080 |
| **Right-Amygdala** | **0.157** | **0.018** | **rh_lateraloccipital_volume** | **0.173** | **0.011** |
| Right-Accumbens-area | 0.026 | 0.363 | **rh_lateralorbitofrontal_volume** | **0.125** | **0.048** |
| Right-VentralDC | 0.106 | 0.080 | rh_lingual_volume | 0.081 | 0.141 |
| CC_Posterior | 0.013 | 0.429 | **rh_medialorbitofrontal_volume** | **0.189** | **0.006** |
| CC_Mid_Posterior | -0.032 | 0.337 | **rh_middletemporal_volume** | **0.349** | **>0.001** |
| CC_Central | 0.083 | 0.134 | **rh_parahippocampal_volume** | **0.178** | **0.009** |
| CC_Mid_Anterior | 0.105 | 0.081 | **rh_paracentral_volume** | **0.167** | **0.013** |
| CC_Anterior | 0.012 | 0.436 | **rh_parsopercularis_volume** | **0.126** | **0.047** |
| **lh_bankssts_volume** | **0.335** | **>0.001** | **rh_parsorbitalis_volume** | **0.139** | **0.032** |
| lh_caudalanteriorcingulate_volume | -0.092 | 0.110 | **rh_parstriangularis_volume** | **0.212** | **0.002** |
| **lh_caudalmiddlefrontal_volume** | **0.240** | **0.001** | rh_pericalcarine_volume | 0.052 | 0.245 |
| lh_cuneus_volume | 0.012 | 0.435 | rh_postcentral_volume | 0.089 | 0.120 |
| **lh_entorhinal_volume** | **0.210** | **0.003** | **rh_posteriorcingulate_volume** | **0.265** | **>0.001** |
| **lh_fusiform_volume** | **0.371** | **>0.001** | rh_precentral_volume | 0.034 | 0.327 |
| **lh_inferiorparietal_volume** | **0.377** | **>0.001** | **rh_precuneus_volume** | **0.237** | **0.001** |
| **lh_inferiortemporal_volume** | **0.227** | **0.001** | rh_rostralanteriorcingulate_volume | 0.057 | 0.226 |
| **lh_isthmuscingulate_volume** | **0.130** | **0.043** | **rh_rostralmiddlefrontal_volume** | **0.271** | **>0.001** |
| lh_lateraloccipital_volume | 0.122 | 0.053 | **rh_superiorfrontal_volume** | **0.249** | **>0.001** |
| **lh_lateralorbitofrontal_volume** | **0.145** | **0.027** | **rh_superiorparietal_volume** | **0.241** | **0.001** |
| lh_lingual_volume | 0.112 | 0.069 | **rh_superiortemporal_volume** | **0.334** | **>0.001** |
| **lh_medialorbitofrontal_volume** | **0.197** | **0.004** | **rh_supramarginal_volume** | **0.230** | **0.001** |
| **lh_middletemporal_volume** | **0.344** | **>0.001** | rh_frontalpole_volume | -0.029 | 0.352 |
| **lh_parahippocampal_volume** | **0.167** | **0.013** | rh_temporalpole_volume | 0.089 | 0.118 |
| **lh_paracentral_volume** | **0.125** | **0.048** | rh_transversetemporal_volume | 0.033 | 0.332 |
| **lh_parsopercularis_volume** | **0.247** | **0.001** | **rh_insula_volume** | **0.204** | **0.003** |
| lh_parsorbitalis_volume | 0.075 | 0.160 | | | |

### 4. Univariate analysis – ADNI dataset – HC vs AD

Table 3 - Statistical significance measured by the Mann-Whitney U test and effect size measured by Cliff's delta absolute value based on the comparison of the reconstruction error for each brain region between the HC and the AD groups from the ADNI dataset. The regions with p-value <= 0.05 are highlighted in bold.

| Regions | Effect size | p-value | Regions | Effect size | p-value |
|---|---|---|---|---|---|
| **Left-Lateral-Ventricle** | **-0.552** | **>0.001** | **lh_parstriangularis_volume** | **0.420** | **>0.001** |
| **Left-Inf-Lat-Vent** | **-0.708** | **>0.001** | **lh_pericalcarine_volume** | **0.246** | **0.001** |
| Left-Cerebellum-White-Matter | -0.040 | 0.316 | **lh_postcentral_volume** | **0.330** | **>0.001** |
| Left-Cerebellum-Cortex | 0.103 | 0.106 | **lh_posteriorcingulate_volume** | **0.414** | **>0.001** |
| **Left-Thalamus-Proper** | **0.488** | **>0.001** | **lh_precentral_volume** | **0.317** | **>0.001** |
| **Left-Caudate** | **0.179** | **0.015** | **lh_precuneus_volume** | **0.640** | **>0.001** |
| **Left-Putamen** | **0.378** | **>0.001** | **lh_rostralanteriorcingulate_volume** | **0.328** | **>0.001** |
| Left-Pallidum | -0.052 | 0.264 | **lh_rostralmiddlefrontal_volume** | **0.581** | **>0.001** |
| **3rd-Ventricle** | **-0.542** | **>0.001** | **lh_superiorfrontal_volume** | **0.633** | **>0.001** |
| 4th-Ventricle | -0.075 | 0.181 | **lh_superiorparietal_volume** | **0.424** | **>0.001** |
| **Brain-Stem** | **0.189** | **0.011** | **lh_superiortemporal_volume** | **0.676** | **>0.001** |
| **Left-Hippocampus** | **0.748** | **>0.001** | **lh_supramarginal_volume** | **0.587** | **>0.001** |
| **Left-Amygdala** | **0.768** | **>0.001** | **lh_frontalpole_volume** | **0.204** | **0.007** |
| **CSF** | **-0.528** | **>0.001** | **lh_temporalpole_volume** | **0.203** | **0.007** |
| **Left-Accumbens-area** | **0.458** | **>0.001** | **lh_transversetemporal_volume** | **0.276** | **>0.001** |
| **Left-VentralDC** | **0.277** | **>0.001** | **lh_insula_volume** | **0.401** | **>0.001** |
| **Right-Lateral-Ventricle** | **-0.533** | **>0.001** | **rh_bankssts_volume** | **0.554** | **>0.001** |
| **Right-Inf-Lat-Vent** | **-0.705** | **>0.001** | rh_caudalanteriorcingulate_volume | 0.082 | 0.160 |
| Right-Cerebellum-White-Matter | -0.080 | 0.166 | **rh_caudalmiddlefrontal_volume** | **0.442** | **>0.001** |
| **Right-Cerebellum-Cortex** | **0.153** | **0.032** | **rh_cuneus_volume** | **0.148** | **0.037** |
| **Right-Thalamus-Proper** | **0.499** | **>0.001** | **rh_entorhinal_volume** | **0.430** | **>0.001** |
| **Right-Caudate** | **0.136** | **0.049** | **rh_fusiform_volume** | **0.698** | **>0.001** |
| **Right-Putamen** | **0.371** | **>0.001** | **rh_inferiorparietal_volume** | **0.618** | **>0.001** |
| Right-Pallidum | 0.002 | 0.491 | **rh_inferiortemporal_volume** | **0.621** | **>0.001** |
| **Right-Hippocampus** | **0.719** | **>0.001** | **rh_isthmuscingulate_volume** | **0.412** | **>0.001** |
| **Right-Amygdala** | **0.685** | **>0.001** | **rh_lateraloccipital_volume** | **0.422** | **>0.001** |
| **Right-Accumbens-area** | **0.349** | **>0.001** | **rh_lateralorbitofrontal_volume** | **0.422** | **>0.001** |
| **Right-VentralDC** | **0.230** | **0.003** | **rh_lingual_volume** | **0.231** | **0.003** |
| CC_Posterior | -0.026 | 0.376 | **rh_medialorbitofrontal_volume** | **0.415** | **>0.001** |
| **CC_Mid_Posterior** | **0.177** | **0.016** | **rh_middletemporal_volume** | **0.697** | **>0.001** |
| **CC_Central** | **0.383** | **>0.001** | **rh_parahippocampal_volume** | **0.416** | **>0.001** |
| **CC_Mid_Anterior** | **0.306** | **>0.001** | **rh_paracentral_volume** | **0.264** | **0.001** |
| CC_Anterior | 0.059 | 0.238 | **rh_parsopercularis_volume** | **0.310** | **>0.001** |
| **lh_bankssts_volume** | **0.656** | **>0.001** | **rh_parsorbitalis_volume** | **0.349** | **>0.001** |
| lh_caudalanteriorcingulate_volume | -0.035 | 0.337 | **rh_parstriangularis_volume** | **0.266** | **0.001** |
| **lh_caudalmiddlefrontal_volume** | **0.487** | **>0.001** | **rh_pericalcarine_volume** | **0.211** | **0.005** |
| **lh_cuneus_volume** | **0.200** | **0.008** | **rh_postcentral_volume** | **0.318** | **>0.001** |
| **lh_entorhinal_volume** | **0.514** | **>0.001** | **rh_posteriorcingulate_volume** | **0.427** | **>0.001** |
| **lh_fusiform_volume** | **0.621** | **>0.001** | **rh_precentral_volume** | **0.280** | **>0.001** |
| **lh_inferiorparietal_volume** | **0.644** | **>0.001** | **rh_precuneus_volume** | **0.573** | **>0.001** |
| **lh_inferiortemporal_volume** | **0.641** | **>0.001** | **rh_rostralanteriorcingulate_volume** | **0.143** | **0.041** |
| **lh_isthmuscingulate_volume** | **0.446** | **>0.001** | **rh_rostralmiddlefrontal_volume** | **0.555** | **>0.001** |
| **lh_lateraloccipital_volume** | **0.388** | **>0.001** | **rh_superiorfrontal_volume** | **0.450** | **>0.001** |
| **lh_lateralorbitofrontal_volume** | **0.439** | **>0.001** | **rh_superiorparietal_volume** | **0.519** | **>0.001** |
| **lh_lingual_volume** | **0.354** | **>0.001** | **rh_superiortemporal_volume** | **0.613** | **>0.001** |
| **lh_medialorbitofrontal_volume** | **0.453** | **>0.001** | **rh_supramarginal_volume** | **0.572** | **>0.001** |
| **lh_middletemporal_volume** | **0.686** | **>0.001** | rh_frontalpole_volume | 0.072 | 0.190 |
| **lh_parahippocampal_volume** | **0.458** | **>0.001** | **rh_temporalpole_volume** | **0.297** | **>0.001** |
| **lh_paracentral_volume** | **0.250** | **0.001** | **rh_transversetemporal_volume** | **0.230** | **0.003** |
| **lh_parsopercularis_volume** | **0.382** | **>0.001** | **rh_insula_volume** | **0.468** | **>0.001** |
| **lh_parsorbitalis_volume** | **0.271** | **0.001** | | | |

#### 5. Univariate analysis – AIBL dataset – HC vs MCI

Table 4 - Statistical significance measured by the Mann-Whitney U test and effect size measured by Cliff’s delta absolute value based on the comparison of the reconstruction error for each brain region between the HC and the MCI groups from the AIBL dataset. The regions with p-value <= 0.05 are highlighted in bold.

| Regions | Effect size | p-value | Regions | Effect size | p-value |
|---|---|---|---|---|---|
| Left-Lateral-Ventricle | -0.125 | 0.088 | lh_parstriangularis_volume | -0.008 | 0.466 |
| **Left-Inf-Lat-Vent** | **-0.273** | **0.002** | lh_pericalcarine_volume | 0.109 | 0.120 |
| Left-Cerebellum-White-Matter | 0.128 | 0.084 | lh_postcentral_volume | 0.144 | 0.059 |
| Left-Cerebellum-Cortex | 0.140 | 0.065 | lh_posteriorcingulate_volume | 0.075 | 0.210 |
| Left-Thalamus-Proper | 0.104 | 0.132 | lh_precentral_volume | 0.067 | 0.236 |
| Left-Caudate | 0.115 | 0.106 | **lh_precuneus_volume** | **0.294** | **0.001** |
| Left-Putamen | -0.013 | 0.444 | lh_rostralanteriorcingulate_volume | 0.090 | 0.164 |
| Left-Pallidum | -0.045 | 0.313 | **lh_rostralmiddlefrontal_volume** | **0.205** | **0.013** |
| **3rd-Ventricle** | **-0.234** | **0.006** | **lh_superiorfrontal_volume** | **0.236** | **0.005** |
| 4th-Ventricle | -0.037 | 0.343 | lh_superiorparietal_volume | 0.077 | 0.202 |
| Brain-Stem | 0.005 | 0.480 | **lh_superiortemporal_volume** | **0.261** | **0.002** |
| **Left-Hippocampus** | **0.299** | **0.001** | lh_supramarginal_volume | 0.105 | 0.129 |
| **Left-Amygdala** | **0.218** | **0.009** | lh_frontalpole_volume | 0.116 | 0.106 |
| **CSF** | **-0.323** | **0.000** | **lh_temporalpole_volume** | **0.258** | **0.003** |
| Left-Accumbens-area | -0.019 | 0.417 | lh_transversetemporal_volume | 0.038 | 0.341 |
| Left-VentralDC | -0.017 | 0.427 | **lh_insula_volume** | **0.223** | **0.008** |
| **Right-Lateral-Ventricle** | **-0.167** | **0.035** | rh_bankssts_volume | 0.108 | 0.123 |
| **Right-Inf-Lat-Vent** | **-0.364** | **>0.001** | rh_caudalanteriorcingulate_volume | 0.000 | 0.500 |
| Right-Cerebellum-White-Matter | 0.145 | 0.058 | rh_caudalmiddlefrontal_volume | 0.150 | 0.053 |
| **Right-Cerebellum-Cortex** | **0.159** | **0.042** | rh_cuneus_volume | 0.108 | 0.122 |
| **Right-Thalamus-Proper** | **0.165** | **0.037** | rh_entorhinal_volume | 0.150 | 0.052 |
| Right-Caudate | 0.029 | 0.379 | **rh_fusiform_volume** | **0.243** | **0.004** |
| Right-Putamen | -0.001 | 0.496 | **rh_inferiorparietal_volume** | **0.173** | **0.031** |
| Right-Pallidum | 0.020 | 0.413 | rh_inferiortemporal_volume | 0.140 | 0.065 |
| **Right-Hippocampus** | **0.311** | **>0.001** | **rh_isthmuscingulate_volume** | **0.260** | **0.003** |
| **Right-Amygdala** | **0.210** | **0.012** | rh_lateraloccipital_volume | 0.129 | 0.082 |
| Right-Accumbens-area | -0.025 | 0.392 | **rh_lateralorbitofrontal_volume** | **0.253** | **0.003** |
| Right-VentralDC | -0.077 | 0.201 | **rh_lingual_volume** | **0.189** | **0.020** |
| CC_Posterior | 0.008 | 0.467 | rh_medialorbitofrontal_volume | 0.115 | 0.106 |
| CC_Mid_Posterior | 0.117 | 0.104 | **rh_middletemporal_volume** | **0.176** | **0.029** |
| CC_Central | 0.048 | 0.302 | **rh_parahippocampal_volume** | **0.154** | **0.048** |
| CC_Mid_Anterior | 0.118 | 0.101 | rh_paracentral_volume | 0.134 | 0.073 |
| CC_Anterior | 0.062 | 0.250 | rh_parsopercularis_volume | 0.106 | 0.126 |
| **lh_bankssts_volume** | **0.203** | **0.014** | rh_parsorbitalis_volume | 0.055 | 0.278 |
| lh_caudalanteriorcingulate_volume | 0.030 | 0.374 | rh_parstriangularis_volume | 0.093 | 0.158 |
| **lh_caudalmiddlefrontal_volume** | **0.189** | **0.021** | rh_pericalcarine_volume | 0.077 | 0.203 |
| **lh_cuneus_volume** | **0.193** | **0.018** | **rh_postcentral_volume** | **0.173** | **0.031** |
| **lh_entorhinal_volume** | **0.340** | **>0.001** | rh_posteriorcingulate_volume | 0.063 | 0.247 |
| **lh_fusiform_volume** | **0.249** | **0.003** | rh_precentral_volume | 0.131 | 0.079 |
| lh_inferiorparietal_volume | 0.089 | 0.168 | **rh_precuneus_volume** | **0.223** | **0.008** |
| lh_inferiortemporal_volume | 0.143 | 0.061 | **rh_rostralanteriorcingulate_volume** | **0.177** | **0.028** |
| lh_isthmuscingulate_volume | 0.136 | 0.071 | rh_rostralmiddlefrontal_volume | 0.091 | 0.162 |
| lh_lateraloccipital_volume | 0.142 | 0.062 | **rh_superiorfrontal_volume** | **0.250** | **0.003** |
| **lh_lateralorbitofrontal_volume** | **0.248** | **0.004** | **rh_superiorparietal_volume** | **0.241** | **0.005** |
| lh_lingual_volume | 0.126 | 0.087 | **rh_superiortemporal_volume** | **0.239** | **0.005** |
| **lh_medialorbitofrontal_volume** | **0.292** | **0.001** | rh_supramarginal_volume | 0.092 | 0.160 |
| lh_middletemporal_volume | 0.120 | 0.097 | rh_frontalpole_volume | 0.081 | 0.189 |
| lh_parahippocampal_volume | 0.095 | 0.151 | **rh_temporalpole_volume** | **0.188** | **0.021** |
| lh_paracentral_volume | 0.101 | 0.138 | **rh_transversetemporal_volume** | **0.197** | **0.017** |
| lh_parsopercularis_volume | 0.049 | 0.298 | **rh_insula_volume** | **0.252** | **0.003** |
| lh_parsorbitalis_volume | 0.038 | 0.341 | | | |

### 6. Univariate analysis – AIBL dataset – HC vs AD

Table 5 - Statistical significance measured by the Mann-Whitney U test and effect size measured by Cliff’s delta absolute value based on the comparison of the reconstruction error for each brain region between the HC and the AD groups from AIBL dataset. The regions with p-value <= 0.05 are highlighted in bold.

| Regions | Effect size | p-value | Regions | Effect size | p-value |
|---|---|---|---|---|---|
| **Left-Lateral-Ventricle** | **-0.584** | **>0.001** | **lh_parstriangularis_volume** | **0.275** | **0.003** |
| **Left-Inf-Lat-Vent** | **-0.803** | **>0.001** | lh_pericalcarine_volume | 0.133 | 0.092 |
| **Left-Cerebellum-White-Matter** | **0.224** | **0.013** | lh_postcentral_volume | 0.137 | 0.086 |
| **Left-Cerebellum-Cortex** | **0.307** | **0.001** | **lh_posteriorcingulate_volume** | **0.416** | **0.000** |
| **Left-Thalamus-Proper** | **0.335** | **>0.001** | lh_precentral_volume | 0.075 | 0.228 |
| Left-Caudate | 0.147 | 0.072 | **lh_precuneus_volume** | **0.721** | **>0.001** |
| **Left-Putamen** | **0.284** | **0.002** | **lh_rostralanteriorcingulate_volume** | **0.387** | **>0.001** |
| Left-Pallidum | 0.151 | 0.066 | **lh_rostralmiddlefrontal_volume** | **0.510** | **>0.001** |
| **3rd-Ventricle** | **-0.510** | **>0.001** | **lh_superiorfrontal_volume** | **0.326** | **0.001** |
| 4th-Ventricle | -0.079 | 0.216 | **lh_superiorparietal_volume** | **0.447** | **>0.001** |
| **Brain-Stem** | **0.197** | **0.025** | **lh_superiortemporal_volume** | **0.481** | **>0.001** |
| **Left-Hippocampus** | **0.729** | **>0.001** | **lh_supramarginal_volume** | **0.382** | **>0.001** |
| **Left-Amygdala** | **0.666** | **>0.001** | **lh_frontalpole_volume** | **0.184** | **0.034** |
| **CSF** | **-0.492** | **>0.001** | **lh_temporalpole_volume** | **0.238** | **0.009** |
| **Left-Accumbens-area** | **0.217** | **0.015** | lh_transversetemporal_volume | 0.113 | 0.130 |
| **Left-VentralDC** | **0.226** | **0.012** | **lh_insula_volume** | **0.272** | **0.003** |
| **Right-Lateral-Ventricle** | **-0.610** | **>0.001** | **rh_bankssts_volume** | **0.540** | **0.000** |
| **Right-Inf-Lat-Vent** | **-0.764** | **>0.001** | **rh_caudalanteriorcingulate_volume** | **0.241** | **0.008** |
| **Right-Cerebellum-White-Matter** | **0.203** | **0.022** | **rh_caudalmiddlefrontal_volume** | **0.386** | **>0.001** |
| **Right-Cerebellum-Cortex** | **0.313** | **0.001** | **rh_cuneus_volume** | **0.287** | **0.002** |
| **Right-Thalamus-Proper** | **0.356** | **>0.001** | **rh_entorhinal_volume** | **0.734** | **>0.001** |
| Right-Caudate | 0.151 | 0.067 | **rh_fusiform_volume** | **0.533** | **>0.001** |
| **Right-Putamen** | **0.276** | **0.003** | **rh_inferiorparietal_volume** | **0.728** | **>0.001** |
| Right-Pallidum | 0.141 | 0.080 | **rh_inferiortemporal_volume** | **0.720** | **>0.001** |
| **Right-Hippocampus** | **0.751** | **>0.001** | **rh_isthmuscingulate_volume** | **0.595** | **>0.001** |
| **Right-Amygdala** | **0.737** | **>0.001** | **rh_lateraloccipital_volume** | **0.485** | **>0.001** |
| **Right-Accumbens-area** | **0.362** | **>0.001** | **rh_lateralorbitofrontal_volume** | **0.407** | **>0.001** |
| **Right-VentralDC** | **0.270** | **0.004** | **rh_lingual_volume** | **0.339** | **>0.001** |
| CC_Posterior | 0.070 | 0.244 | **rh_medialorbitofrontal_volume** | **0.340** | **>0.001** |
| **CC_Mid_Posterior** | **0.307** | **0.001** | **rh_middletemporal_volume** | **0.609** | **>0.001** |
| **CC_Central** | **0.393** | **>0.001** | **rh_parahippocampal_volume** | **0.576** | **>0.001** |
| **CC_Mid_Anterior** | **0.427** | **>0.001** | rh_paracentral_volume | 0.103 | 0.153 |
| CC_Anterior | 0.084 | 0.203 | **rh_parsopercularis_volume** | **0.305** | **0.001** |
| **lh_bankssts_volume** | **0.389** | **>0.001** | **rh_parsorbitalis_volume** | **0.326** | **0.001** |
| lh_caudalanteriorcingulate_volume | 0.045 | 0.326 | **rh_parstriangularis_volume** | **0.309** | **0.001** |
| **lh_caudalmiddlefrontal_volume** | **0.312** | **0.001** | rh_pericalcarine_volume | 0.161 | 0.054 |
| **lh_cuneus_volume** | **0.357** | **>0.001** | rh_postcentral_volume | 0.046 | 0.322 |
| **lh_entorhinal_volume** | **0.719** | **>0.001** | **rh_posteriorcingulate_volume** | **0.360** | **>0.001** |
| **lh_fusiform_volume** | **0.586** | **>0.001** | rh_precentral_volume | 0.074 | 0.231 |
| **lh_inferiorparietal_volume** | **0.636** | **>0.001** | **rh_precuneus_volume** | **0.652** | **>0.001** |
| **lh_inferiortemporal_volume** | **0.585** | **>0.001** | **rh_rostralanteriorcingulate_volume** | **0.388** | **>0.001** |
| **lh_isthmuscingulate_volume** | **0.590** | **>0.001** | **rh_rostralmiddlefrontal_volume** | **0.469** | **>0.001** |
| **lh_lateraloccipital_volume** | **0.589** | **>0.001** | **rh_superiorfrontal_volume** | **0.304** | **0.001** |
| **lh_lateralorbitofrontal_volume** | **0.407** | **>0.001** | **rh_superiorparietal_volume** | **0.596** | **>0.001** |
| **lh_lingual_volume** | **0.283** | **0.002** | **rh_superiortemporal_volume** | **0.441** | **>0.001** |
| **lh_medialorbitofrontal_volume** | **0.214** | **0.016** | **rh_supramarginal_volume** | **0.496** | **>0.001** |
| **lh_middletemporal_volume** | **0.624** | **>0.001** | rh_frontalpole_volume | 0.092 | 0.179 |
| **lh_parahippocampal_volume** | **0.485** | **>0.001** | **rh_temporalpole_volume** | **0.381** | **>0.001** |
| lh_paracentral_volume | 0.137 | 0.087 | **rh_transversetemporal_volume** | **0.219** | **0.014** |
| **lh_parsopercularis_volume** | **0.365** | **>0.001** | **rh_insula_volume** | **0.439** | **>0.001** |
| **lh_parsorbitalis_volume** | **0.256** | **0.005** | | | |

### 7. Univariate analysis – ARWIBO dataset – HC vs MCI

Table 6 - Statistical significance measured by the Mann-Whitney U test and effect size measured by Cliff's delta absolute value based on the comparison of the reconstruction error for each brain region between the HC and the MCI groups from the ARWIBO dataset. The regions with p-value <= 0.05 are highlighted in bold.

| Regions | Effect size | p-value | Regions | Effect size | p-value |
|---|---|---|---|---|---|
| **Left-Lateral-Ventricle** | **-0.251** | **0.001** | **lh_parstriangularis_volume** | **0.199** | **0.007** |
| **Left-Inf-Lat-Vent** | **-0.250** | **0.001** | lh_pericalcarine_volume | -0.018 | 0.412 |
| **Left-Cerebellum-White-Matter** | **0.167** | **0.020** | lh_postcentral_volume | 0.043 | 0.296 |
| Left-Cerebellum-Cortex | -0.085 | 0.147 | **lh_posteriorcingulate_volume** | **0.236** | **0.002** |
| **Left-Thalamus-Proper** | **0.210** | **0.005** | **lh_precentral_volume** | **0.145** | **0.036** |
| Left-Caudate | 0.047 | 0.280 | **lh_precuneus_volume** | **0.229** | **0.002** |
| Left-Putamen | 0.084 | 0.149 | **lh_rostralanteriorcingulate_volume** | **0.139** | **0.043** |
| Left-Pallidum | 0.010 | 0.452 | **lh_rostralmiddlefrontal_volume** | **0.189** | **0.010** |
| **3rd-Ventricle** | **-0.235** | **0.002** | **lh_superiorfrontal_volume** | **0.226** | **0.003** |
| **4th-Ventricle** | **-0.170** | **0.018** | lh_superiorparietal_volume | 0.132 | 0.051 |
| **Brain-Stem** | **0.148** | **0.034** | lh_superiortemporal_volume | -0.020 | 0.402 |
| **Left-Hippocampus** | **0.276** | **>0.001** | lh_supramarginal_volume | 0.070 | 0.193 |
| Left-Amygdala | 0.122 | 0.066 | lh_frontalpole_volume | -0.037 | 0.323 |
| **CSF** | **-0.285** | **>0.001** | lh_temporalpole_volume | 0.071 | 0.190 |
| **Left-Accumbens-area** | **0.277** | **>0.001** | lh_transversetemporal_volume | -0.132 | 0.051 |
| Left-VentralDC | 0.119 | 0.071 | **lh_insula_volume** | **0.185** | **0.011** |
| **Right-Lateral-Ventricle** | **-0.261** | **0.001** | **rh_bankssts_volume** | **0.247** | **0.001** |
| **Right-Inf-Lat-Vent** | **-0.335** | **>0.001** | rh_caudalanteriorcingulate_volume | 0.039 | 0.315 |
| Right-Cerebellum-White-Matter | 0.037 | 0.325 | **rh_caudalmiddlefrontal_volume** | **0.243** | **0.001** |
| Right-Cerebellum-Cortex | -0.048 | 0.276 | rh_cuneus_volume | -0.026 | 0.376 |
| **Right-Thalamus-Proper** | **0.170** | **0.018** | **rh_entorhinal_volume** | **0.272** | **>0.001** |
| Right-Caudate | 0.110 | 0.087 | **rh_fusiform_volume** | **0.306** | **>0.001** |
| Right-Putamen | 0.095 | 0.121 | **rh_inferiorparietal_volume** | **0.271** | **>0.001** |
| Right-Pallidum | 0.085 | 0.148 | **rh_inferiortemporal_volume** | **0.344** | **>0.001** |
| **Right-Hippocampus** | **0.269** | **>0.001** | **rh_isthmuscingulate_volume** | **0.137** | **0.045** |
| **Right-Amygdala** | **0.237** | **0.002** | **rh_lateraloccipital_volume** | **0.135** | **0.048** |
| Right-Accumbens-area | 0.109 | 0.088 | **rh_lateralorbitofrontal_volume** | **0.195** | **0.008** |
| Right-VentralDC | 0.106 | 0.095 | **rh_lingual_volume** | **0.165** | **0.021** |
| CC_Posterior | 0.110 | 0.088 | **rh_medialorbitofrontal_volume** | **0.232** | **0.002** |
| CC_Mid_Posterior | 0.120 | 0.069 | **rh_middletemporal_volume** | **0.228** | **0.002** |
| **CC_Central** | **0.147** | **0.035** | **rh_parahippocampal_volume** | **0.228** | **0.002** |
| CC_Mid_Anterior | 0.089 | 0.135 | **rh_paracentral_volume** | **0.166** | **0.020** |
| CC_Anterior | 0.068 | 0.202 | rh_parsopercularis_volume | 0.098 | 0.114 |
| lh_bankssts_volume | 0.085 | 0.146 | **rh_parsorbitalis_volume** | **0.181** | **0.013** |
| lh_caudalanteriorcingulate_volume | 0.125 | 0.062 | **rh_parstriangularis_volume** | **0.168** | **0.019** |
| **lh_caudalmiddlefrontal_volume** | **0.147** | **0.035** | rh_pericalcarine_volume | 0.093 | 0.126 |
| lh_cuneus_volume | -0.025 | 0.378 | rh_postcentral_volume | -0.020 | 0.400 |
| lh_entorhinal_volume | 0.203 | 0.006 | **rh_posteriorcingulate_volume** | **0.244** | **0.001** |
| **lh_fusiform_volume** | **0.262** | **0.001** | **rh_precentral_volume** | **0.198** | **0.007** |
| lh_inferiorparietal_volume | 0.198 | 0.007 | **rh_precuneus_volume** | **0.256** | **0.001** |
| **lh_inferiortemporal_volume** | **0.275** | **>0.001** | **rh_rostralanteriorcingulate_volume** | **0.191** | **0.009** |
| **lh_isthmuscingulate_volume** | **0.155** | **0.028** | rh_rostralmiddlefrontal_volume | 0.119 | 0.070 |
| **lh_lateraloccipital_volume** | **0.176** | **0.015** | **rh_superiorfrontal_volume** | **0.261** | **0.001** |
| **lh_lateralorbitofrontal_volume** | **0.193** | **0.009** | **rh_superiorparietal_volume** | **0.204** | **0.006** |
| **lh_lingual_volume** | **0.175** | **0.015** | rh_superiortemporal_volume | 0.097 | 0.117 |
| **lh_medialorbitofrontal_volume** | **0.253** | **0.001** | rh_supramarginal_volume | 0.131 | 0.053 |
| **lh_middletemporal_volume** | **0.280** | **>0.001** | rh_frontalpole_volume | -0.023 | 0.389 |
| **lh_parahippocampal_volume** | **0.325** | **>0.001** | rh_temporalpole_volume | 0.084 | 0.150 |
| lh_paracentral_volume | 0.086 | 0.145 | rh_transversetemporal_volume | 0.048 | 0.276 |
| **lh_parsopercularis_volume** | **0.163** | **0.022** | **rh_insula_volume** | **0.149** | **0.033** |
| **lh_parsorbitalis_volume** | **0.209** | **0.005** | | | |

### 8. Univariate analysis – ARWIBO dataset – HC vs AD

Table 7 - Statistical significance measured by the Mann-Whitney U test and effect size measured by Cliff's delta absolute value based on the comparison of the reconstruction error for each brain region between the HC and the AD groups from ARWIBO dataset. The regions with p-value <= 0.05 are highlighted in bold.

| Regions | Effect size | p-value | Regions | Effect size | p-value |
|---|---|---|---|---|---|
| **Left-Lateral-Ventricle** | **-0.547** | **>0.001** | **lh_parstriangularis_volume** | **0.233** | **0.012** |
| **Left-Inf-Lat-Vent** | **-0.758** | **>0.001** | lh_pericalcarine_volume | 0.080 | 0.220 |
| Left-Cerebellum-White-Matter | 0.063 | 0.272 | lh_postcentral_volume | -0.123 | 0.117 |
| Left-Cerebellum-Cortex | -0.017 | 0.435 | **lh_posteriorcingulate_volume** | **0.435** | **>0.001** |
| **Left-Thalamus-Proper** | **0.282** | **0.003** | lh_precentral_volume | -0.029 | 0.391 |
| **Left-Caudate** | **0.321** | **0.001** | **lh_precuneus_volume** | **0.599** | **>0.001** |
| **Left-Putamen** | **0.287** | **0.003** | **lh_rostralanteriorcingulate_volume** | **0.289** | **0.003** |
| Left-Pallidum | -0.156 | 0.065 | **lh_rostralmiddlefrontal_volume** | **0.373** | **>0.001** |
| **3rd-Ventricle** | **-0.517** | **>0.001** | **lh_superiorfrontal_volume** | **0.362** | **>0.001** |
| 4th-Ventricle | -0.050 | 0.314 | **lh_superiorparietal_volume** | **0.303** | **0.002** |
| Brain-Stem | 0.126 | 0.112 | **lh_superiortemporal_volume** | **0.382** | **>0.001** |
| **Left-Hippocampus** | **0.790** | **>0.001** | **lh_supramarginal_volume** | **0.342** | **>0.001** |
| **Left-Amygdala** | **0.649** | **>0.001** | **lh_frontalpole_volume** | **0.211** | **0.020** |
| **CSF** | **-0.518** | **>0.001** | **lh_temporalpole_volume** | **0.477** | **>0.001** |
| **Left-Accumbens-area** | **0.325** | **0.001** | **lh_transversetemporal_volume** | **0.216** | **0.018** |
| Left-VentralDC | 0.138 | 0.091 | **lh_insula_volume** | **0.510** | **>0.001** |
| **Right-Lateral-Ventricle** | **-0.478** | **>0.001** | **rh_bankssts_volume** | **0.486** | **>0.001** |
| **Right-Inf-Lat-Vent** | **-0.768** | **>0.001** | rh_caudalanteriorcingulate_volume | 0.042 | 0.342 |
| Right-Cerebellum-White-Matter | 0.098 | 0.170 | rh_caudalmiddlefrontal_volume | 0.129 | 0.105 |
| Right-Cerebellum-Cortex | -0.024 | 0.409 | rh_cuneus_volume | -0.009 | 0.467 |
| Right-Thalamus-Proper | 0.117 | 0.129 | **rh_entorhinal_volume** | **0.603** | **>0.001** |
| **Right-Caudate** | **0.301** | **0.002** | **rh_fusiform_volume** | **0.506** | **>0.001** |
| **Right-Putamen** | **0.272** | **0.004** | **rh_inferiorparietal_volume** | **0.599** | **>0.001** |
| Right-Pallidum | -0.042 | 0.344 | **rh_inferiortemporal_volume** | **0.415** | **>0.001** |
| **Right-Hippocampus** | **0.635** | **>0.001** | **rh_isthmuscingulate_volume** | **0.370** | **>0.001** |
| **Right-Amygdala** | **0.651** | **>0.001** | **rh_lateraloccipital_volume** | **0.397** | **>0.001** |
| **Right-Accumbens-area** | **0.309** | **0.001** | **rh_lateralorbitofrontal_volume** | **0.305** | **0.002** |
| Right-VentralDC | 0.093 | 0.184 | **rh_lingual_volume** | **0.331** | **0.001** |
| CC_Posterior | 0.155 | 0.066 | **rh_medialorbitofrontal_volume** | **0.303** | **0.002** |
| CC_Mid_Posterior | 0.008 | 0.469 | **rh_middletemporal_volume** | **0.519** | **>0.001** |
| **CC_Central** | **0.258** | **0.006** | **rh_parahippocampal_volume** | **0.326** | **0.001** |
| **CC_Mid_Anterior** | **0.289** | **0.002** | rh_paracentral_volume | 0.114 | 0.135 |
| CC_Anterior | 0.041 | 0.345 | rh_parsopercularis_volume | 0.088 | 0.197 |
| **lh_bankssts_volume** | **0.394** | **0.000** | **rh_parsorbitalis_volume** | **0.325** | **0.001** |
| lh_caudalanteriorcingulate_volume | 0.037 | 0.361 | **rh_parstriangularis_volume** | **0.201** | **0.026** |
| **lh_caudalmiddlefrontal_volume** | **0.352** | **>0.001** | rh_pericalcarine_volume | 0.061 | 0.276 |
| lh_cuneus_volume | 0.037 | 0.361 | rh_postcentral_volume | -0.147 | 0.077 |
| **lh_entorhinal_volume** | **0.633** | **>0.001** | **rh_posteriorcingulate_volume** | **0.340** | **>0.001** |
| **lh_fusiform_volume** | **0.657** | **>0.001** | rh_precentral_volume | -0.022 | 0.417 |
| **lh_inferiorparietal_volume** | **0.526** | **>0.001** | **rh_precuneus_volume** | **0.570** | **>0.001** |
| **lh_inferiortemporal_volume** | **0.553** | **>0.001** | **rh_rostralanteriorcingulate_volume** | **0.240** | **0.010** |
| **lh_isthmuscingulate_volume** | **0.344** | **>0.001** | **rh_rostralmiddlefrontal_volume** | **0.304** | **0.002** |
| **lh_lateraloccipital_volume** | **0.322** | **0.001** | **rh_superiorfrontal_volume** | **0.330** | **0.001** |
| **lh_lateralorbitofrontal_volume** | **0.325** | **0.001** | **rh_superiorparietal_volume** | **0.410** | **>0.001** |
| **lh_lingual_volume** | **0.302** | **0.002** | **rh_superiortemporal_volume** | **0.350** | **>0.001** |
| **lh_medialorbitofrontal_volume** | **0.205** | **0.024** | **rh_supramarginal_volume** | **0.380** | **>0.001** |
| **lh_middletemporal_volume** | **0.543** | **>0.001** | rh_frontalpole_volume | 0.136 | 0.094 |
| **lh_parahippocampal_volume** | **0.501** | **>0.001** | **rh_temporalpole_volume** | **0.365** | **>0.001** |
| lh_paracentral_volume | -0.002 | 0.493 | rh_transversetemporal_volume | 0.141 | 0.085 |
| **lh_parsopercularis_volume** | **0.259** | **0.006** | **rh_insula_volume** | **0.426** | **>0.001** |
| **lh_parsorbitalis_volume** | **0.422** | **>0.001** | | | |

#### 9. Univariate analysis – OASIS-1 dataset – HC vs AD

Table 8 - Statistical significance measured by the Mann-Whitney U test and effect size measured by Cliff's delta absolute value based on the comparison of the reconstruction error for each brain region between the HC and the AD groups from the OASIS-1 dataset. The regions with p-value <= 0.05 are highlighted in bold.

| Regions | Effect size | p-value | Regions | Effect size | p-value |
|---|---|---|---|---|---|
| **Left-Lateral-Ventricle** | **-0.469** | **>0.001** | lh_parstriangularis_volume | 0.118 | 0.187 |
| **Left-Inf-Lat-Vent** | **-0.735** | **>0.001** | lh_pericalcarine_volume | -0.010 | 0.472 |
| **Left-Cerebellum-White-Matter** | **0.218** | **0.049** | **lh_postcentral_volume** | **0.362** | **0.003** |
| **Left-Cerebellum-Cortex** | **0.287** | **0.015** | **lh_posteriorcingulate_volume** | **0.359** | **0.003** |
| **Left-Thalamus-Proper** | **0.492** | **>0.001** | **lh_precentral_volume** | **0.330** | **0.006** |
| Left-Caudate | 0.040 | 0.382 | **lh_precuneus_volume** | **0.434** | **0.000** |
| Left-Putamen | 0.210 | 0.056 | lh_rostralanteriorcingulate_volume | 0.102 | 0.221 |
| Left-Pallidum | 0.030 | 0.413 | **lh_rostralmiddlefrontal_volume** | **0.337** | **0.005** |
| 3rd-Ventricle | -0.457 | **>0.001** | **lh_superiorfrontal_volume** | **0.358** | **0.003** |
| 4th-Ventricle | 0.006 | 0.484 | **lh_superiorparietal_volume** | **0.313** | **0.009** |
| **Brain-Stem** | **0.334** | **0.006** | **lh_superiortemporal_volume** | **0.502** | **>0.001** |
| **Left-Hippocampus** | **0.494** | **>0.001** | **lh_supramarginal_volume** | **0.502** | **>0.001** |
| **Left-Amygdala** | **0.481** | **>0.001** | lh_frontalpole_volume | 0.151 | 0.127 |
| **CSF** | **-0.350** | **0.004** | lh_temporalpole_volume | 0.032 | 0.405 |
| **Left-Accumbens-area** | **0.375** | **0.002** | **lh_transversetemporal_volume** | **0.343** | **0.005** |
| **Left-VentralDC** | **0.275** | **0.019** | **lh_insula_volume** | **0.313** | **0.009** |
| **Right-Lateral-Ventricle** | **-0.516** | **>0.001** | **rh_bankssts_volume** | **0.644** | **>0.001** |
| **Right-Inf-Lat-Vent** | **-0.780** | **>0.001** | rh_caudalanteriorcingulate_volume | 0.027 | 0.421 |
| Right-Cerebellum-White-Matter | 0.150 | 0.129 | **rh_caudalmiddlefrontal_volume** | **0.387** | **0.002** |
| **Right-Cerebellum-Cortex** | **0.227** | **0.043** | rh_cuneus_volume | 0.151 | 0.127 |
| **Right-Thalamus-Proper** | **0.521** | **>0.001** | **rh_entorhinal_volume** | **0.566** | **>0.001** |
| Right-Caudate | 0.002 | 0.496 | **rh_fusiform_volume** | **0.511** | **>0.001** |
| **Right-Putamen** | **0.339** | **0.005** | **rh_inferiorparietal_volume** | **0.462** | **>0.001** |
| Right-Pallidum | 0.007 | 0.480 | **rh_inferiortemporal_volume** | **0.572** | **>0.001** |
| **Right-Hippocampus** | **0.623** | **>0.001** | **rh_isthmuscingulate_volume** | **0.274** | **0.019** |
| **Right-Amygdala** | **0.541** | **>0.001** | **rh_lateraloccipital_volume** | **0.249** | **0.030** |
| **Right-Accumbens-area** | **0.424** | **0.001** | **rh_lateralorbitofrontal_volume** | **0.243** | **0.033** |
| **Right-VentralDC** | **0.354** | **0.004** | **rh_lingual_volume** | **0.313** | **0.009** |
| CC_Posterior | 0.183 | 0.084 | **rh_medialorbitofrontal_volume** | **0.312** | **0.009** |
| **CC_Mid_Posterior** | **0.496** | **>0.001** | **rh_middletemporal_volume** | **0.499** | **>0.001** |
| **CC_Central** | **0.503** | **>0.001** | **rh_parahippocampal_volume** | **0.430** | **0.001** |
| **CC_Mid_Anterior** | **0.432** | **0.001** | **rh_paracentral_volume** | **0.234** | **0.038** |
| **CC_Anterior** | **0.375** | **0.002** | **rh_parsopercularis_volume** | **0.297** | **0.012** |
| **lh_bankssts_volume** | **0.428** | **0.001** | **rh_parsorbitalis_volume** | **0.283** | **0.016** |
| lh_caudalanteriorcingulate_volume | 0.011 | 0.468 | **rh_parstriangularis_volume** | **0.283** | **0.016** |
| lh_caudalmiddlefrontal_volume | 0.113 | 0.198 | rh_pericalcarine_volume | 0.074 | 0.288 |
| lh_cuneus_volume | 0.200 | 0.065 | **rh_postcentral_volume** | **0.379** | **0.002** |
| **lh_entorhinal_volume** | **0.339** | **0.005** | rh_posteriorcingulate_volume | 0.183 | 0.084 |
| **lh_fusiform_volume** | **0.375** | **0.002** | **rh_precentral_volume** | **0.428** | **0.001** |
| **lh_inferiorparietal_volume** | **0.452** | **>0.001** | **rh_precuneus_volume** | **0.368** | **0.003** |
| **lh_inferiortemporal_volume** | **0.456** | **>0.001** | rh_rostralanteriorcingulate_volume | 0.114 | 0.195 |
| **lh_isthmuscingulate_volume** | **0.444** | **>0.001** | **rh_rostralmiddlefrontal_volume** | **0.425** | **0.001** |
| **lh_lateraloccipital_volume** | **0.314** | **0.009** | **rh_superiorfrontal_volume** | **0.425** | **0.001** |
| **lh_lateralorbitofrontal_volume** | **0.354** | **0.004** | **rh_superiorparietal_volume** | **0.272** | **0.020** |
| **lh_lingual_volume** | **0.238** | **0.036** | **rh_superiortemporal_volume** | **0.548** | **>0.001** |
| lh_medialorbitofrontal_volume | 0.064 | 0.316 | **rh_supramarginal_volume** | **0.519** | **>0.001** |
| **lh_middletemporal_volume** | **0.507** | **>0.001** | rh_frontalpole_volume | 0.126 | 0.171 |
| **lh_parahippocampal_volume** | **0.458** | **>0.001** | rh_temporalpole_volume | 0.007 | 0.480 |
| **lh_paracentral_volume** | **0.333** | **0.006** | **rh_transversetemporal_volume** | **0.413** | **0.001** |
| **lh_parsopercularis_volume** | **0.245** | **0.032** | **rh_insula_volume** | **0.292** | **0.014** |
| **lh_parsorbitalis_volume** | **0.270** | **0.021** | | | |

### 10. Univariate analysis –MIRIAD dataset – HC vs AD

Table 9 - Statistical significance measured by the Mann-Whitney U test and effect size measured by Cliff’s delta absolute value based on the comparison of the reconstruction error for each brain region between the HC and the AD groups from the MIRIAD dataset. The regions with p-value <= 0.05 are highlighted in bold.

| Regions | Effect size | p-value | Regions | Effect size | p-value |
|---|---|---|---|---|---|
| **Left-Lateral-Ventricle** | **-0.630** | **>0.001** | lh_parstriangularis_volume | 0.034 | 0.361 |
| **Left-Inf-Lat-Vent** | **-0.964** | **>0.001** | lh_pericalcarine_volume | -0.073 | 0.222 |
| Left-Cerebellum-White-Matter | 0.022 | 0.411 | **lh_postcentral_volume** | **0.394** | **>0.001** |
| **Left-Cerebellum-Cortex** | **0.396** | **>0.001** | **lh_posteriorcingulate_volume** | **0.545** | **>0.001** |
| **Left-Thalamus-Proper** | **0.470** | **>0.001** | **lh_precentral_volume** | **0.430** | **>0.001** |
| **Left-Caudate** | **0.338** | **>0.001** | **lh_precuneus_volume** | **0.841** | **>0.001** |
| **Left-Putamen** | **0.386** | **>0.001** | **lh_rostralanteriorcingulate_volume** | **0.440** | **>0.001** |
| Left-Pallidum | 0.092 | 0.169 | **lh_rostralmiddlefrontal_volume** | **0.836** | **>0.001** |
| **3rd-Ventricle** | **-0.385** | **>0.001** | **lh_superiorfrontal_volume** | **0.856** | **>0.001** |
| 4th-Ventricle | 0.059 | 0.269 | **lh_superiorparietal_volume** | **0.741** | **>0.001** |
| **Brain-Stem** | **0.334** | **>0.001** | **lh_superiortemporal_volume** | **0.753** | **>0.001** |
| **Left-Hippocampus** | **0.780** | **>0.001** | **lh_supramarginal_volume** | **0.788** | **>0.001** |
| **Left-Amygdala** | **0.878** | **>0.001** | **lh_frontalpole_volume** | **0.180** | **0.030** |
| **CSF** | **-0.673** | **>0.001** | **lh_temporalpole_volume** | **0.367** | **>0.001** |
| **Left-Accumbens-area** | **0.526** | **>0.001** | **lh_transversetemporal_volume** | **0.532** | **>0.001** |
| **Left-VentralDC** | **0.315** | **>0.001** | **lh_insula_volume** | **0.542** | **>0.001** |
| **Right-Lateral-Ventricle** | **-0.723** | **>0.001** | **rh_bankssts_volume** | **0.641** | **>0.001** |
| **Right-Inf-Lat-Vent** | **-0.932** | **>0.001** | **rh_caudalanteriorcingulate_volume** | **0.374** | **>0.001** |
| Right-Cerebellum-White-Matter | 0.131 | 0.086 | **rh_caudalmiddlefrontal_volume** | **0.471** | **>0.001** |
| **Right-Cerebellum-Cortex** | **0.418** | **>0.001** | **rh_cuneus_volume** | **0.430** | **>0.001** |
| **Right-Thalamus-Proper** | **0.416** | **>0.001** | **rh_entorhinal_volume** | **0.593** | **>0.001** |
| **Right-Caudate** | **0.183** | **0.028** | **rh_fusiform_volume** | **0.819** | **>0.001** |
| **Right-Putamen** | **0.406** | **>0.001** | **rh_inferiorparietal_volume** | **0.807** | **>0.001** |
| Right-Pallidum | 0.060 | 0.265 | **rh_inferiortemporal_volume** | **0.804** | **>0.001** |
| **Right-Hippocampus** | **0.802** | **>0.001** | **rh_isthmuscingulate_volume** | **0.712** | **>0.001** |
| **Right-Amygdala** | **0.877** | **>0.001** | **rh_lateraloccipital_volume** | **0.457** | **>0.001** |
| **Right-Accumbens-area** | **0.558** | **>0.001** | **rh_lateralorbitofrontal_volume** | **0.547** | **>0.001** |
| **Right-VentralDC** | **0.422** | **>0.001** | **rh_lingual_volume** | **0.396** | **>0.001** |
| CC_Posterior | 0.110 | 0.125 | **rh_medialorbitofrontal_volume** | **0.481** | **>0.001** |
| CC_Mid_Posterior | -0.067 | 0.244 | **rh_middletemporal_volume** | **0.825** | **>0.001** |
| **CC_Central** | **0.411** | **>0.001** | **rh_parahippocampal_volume** | **0.488** | **>0.001** |
| **CC_Mid_Anterior** | **0.500** | **>0.001** | rh_paracentral_volume | 0.150 | 0.058 |
| CC_Anterior | 0.083 | 0.193 | **rh_parsopercularis_volume** | **0.363** | **>0.001** |
| **lh_bankssts_volume** | **0.791** | **>0.001** | **rh_parsorbitalis_volume** | **0.232** | **0.008** |
| lh_caudalanteriorcingulate_volume | -0.095 | 0.162 | rh_parstriangularis_volume | -0.110 | 0.126 |
| **lh_caudalmiddlefrontal_volume** | **0.643** | **>0.001** | rh_pericalcarine_volume | 0.046 | 0.315 |
| lh_cuneus_volume | 0.156 | 0.052 | **rh_postcentral_volume** | **0.238** | **0.006** |
| **lh_entorhinal_volume** | **0.446** | **>0.001** | **rh_posteriorcingulate_volume** | **0.575** | **>0.001** |
| **lh_fusiform_volume** | **0.818** | **>0.001** | **rh_precentral_volume** | **0.424** | **>0.001** |
| **lh_inferiorparietal_volume** | **0.869** | **>0.001** | **rh_precuneus_volume** | **0.913** | **>0.001** |
| **lh_inferiortemporal_volume** | **0.884** | **>0.001** | **rh_rostralanteriorcingulate_volume** | **0.245** | **0.005** |
| **lh_isthmuscingulate_volume** | **0.604** | **>0.001** | **rh_rostralmiddlefrontal_volume** | **0.804** | **>0.001** |
| **lh_lateraloccipital_volume** | **0.628** | **>0.001** | **rh_superiorfrontal_volume** | **0.686** | **>0.001** |
| **lh_lateralorbitofrontal_volume** | **0.601** | **>0.001** | **rh_superiorparietal_volume** | **0.812** | **>0.001** |
| **lh_lingual_volume** | **0.304** | **0.001** | **rh_superiortemporal_volume** | **0.645** | **>0.001** |
| **lh_medialorbitofrontal_volume** | **0.535** | **>0.001** | **rh_supramarginal_volume** | **0.857** | **>0.001** |
| **lh_middletemporal_volume** | **0.882** | **>0.001** | **rh_frontalpole_volume** | **0.216** | **0.012** |
| lh_parahippocampal_volume | 0.140 | 0.072 | **rh_temporalpole_volume** | **0.437** | **>0.001** |
| **lh_paracentral_volume** | **0.381** | **>0.001** | **rh_transversetemporal_volume** | **0.393** | **>0.001** |
| **lh_parsopercularis_volume** | **0.314** | **0.001** | **rh_insula_volume** | **0.717** | **>0.001** |
| **lh_parsorbitalis_volume** | **0.333** | **>0.001** | | | |

### 11. Region importance – ADNI dataset – HC vs EMCI

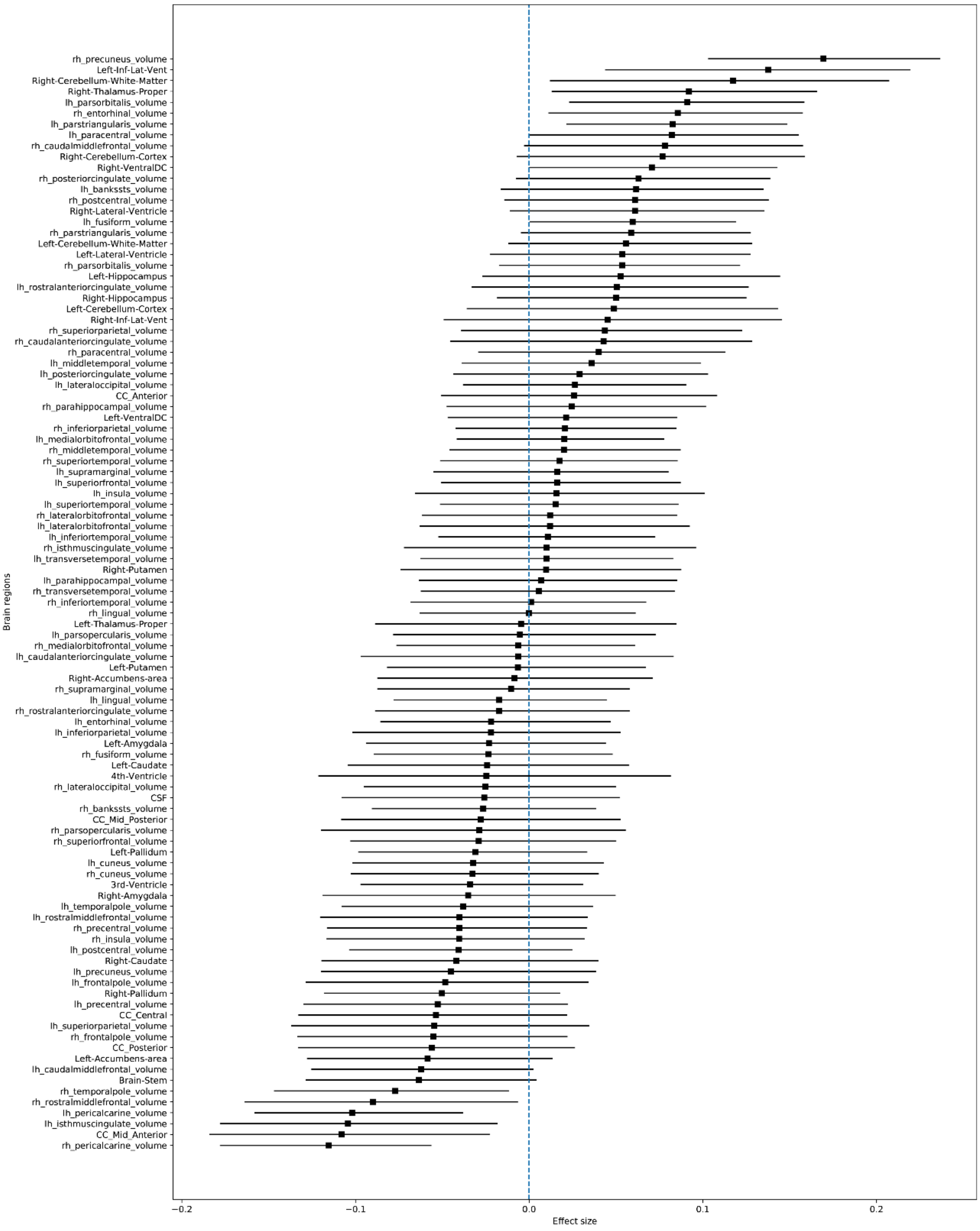


Figure 1 - Regional deviations of the EMCI group from the ADNI dataset. The marker indicates the mean effect size between the HC and the EMCI groups. The horizontal bars indicate the 95% confidence interval calculated using the percentile method on the bootstrap analysis.

### 12. Region importance – ADNI dataset – HC vs LMCI

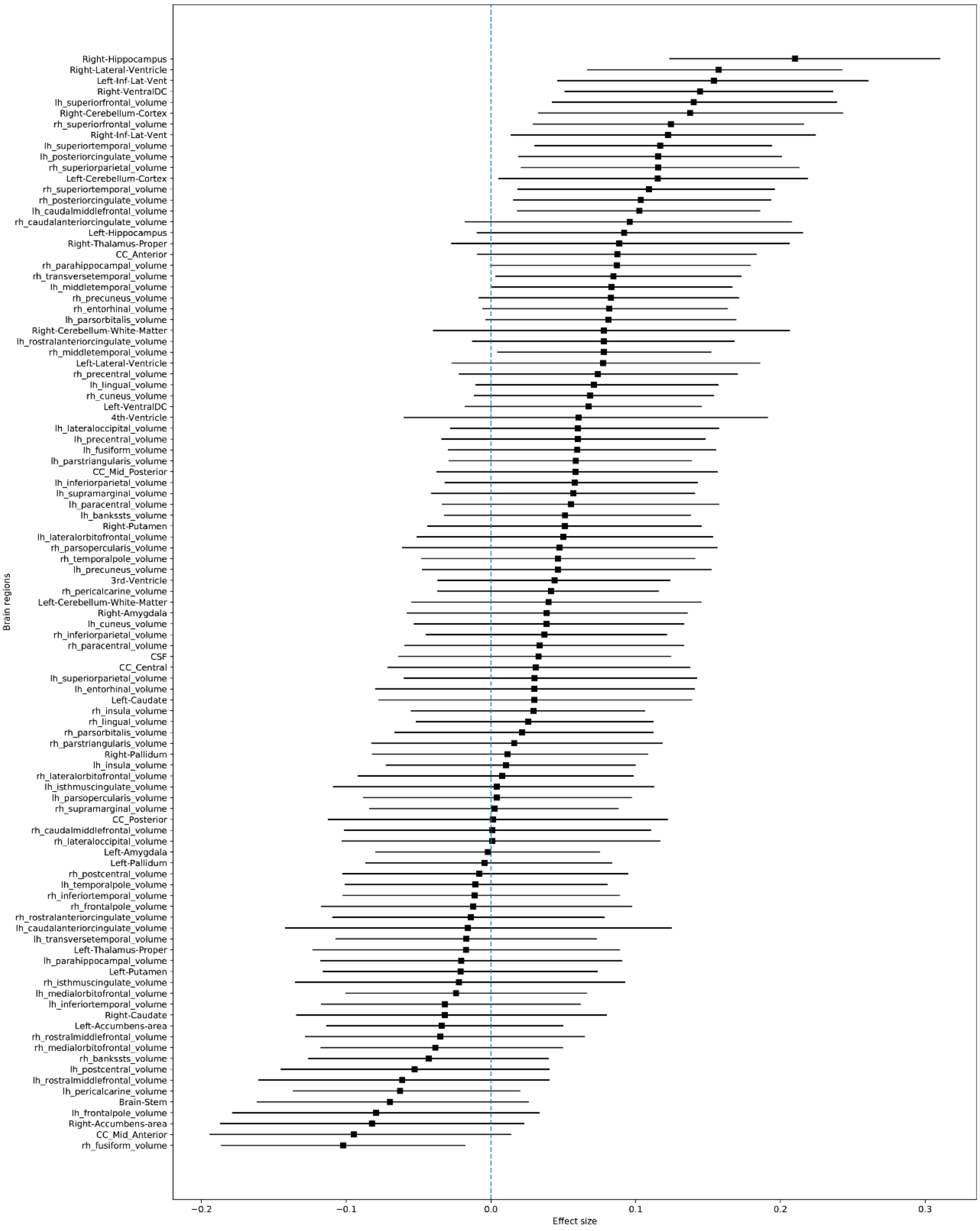


Figure 2 – Regional deviations of the LMCI group from the ADNI dataset. The marker indicates the mean effect size between the HC and the LMCI groups. The horizontal bars indicate the 95% confidence interval calculated using the percentile method on the bootstrap analysis.

#### 13. Region importance – ADNI dataset – HC vs AD

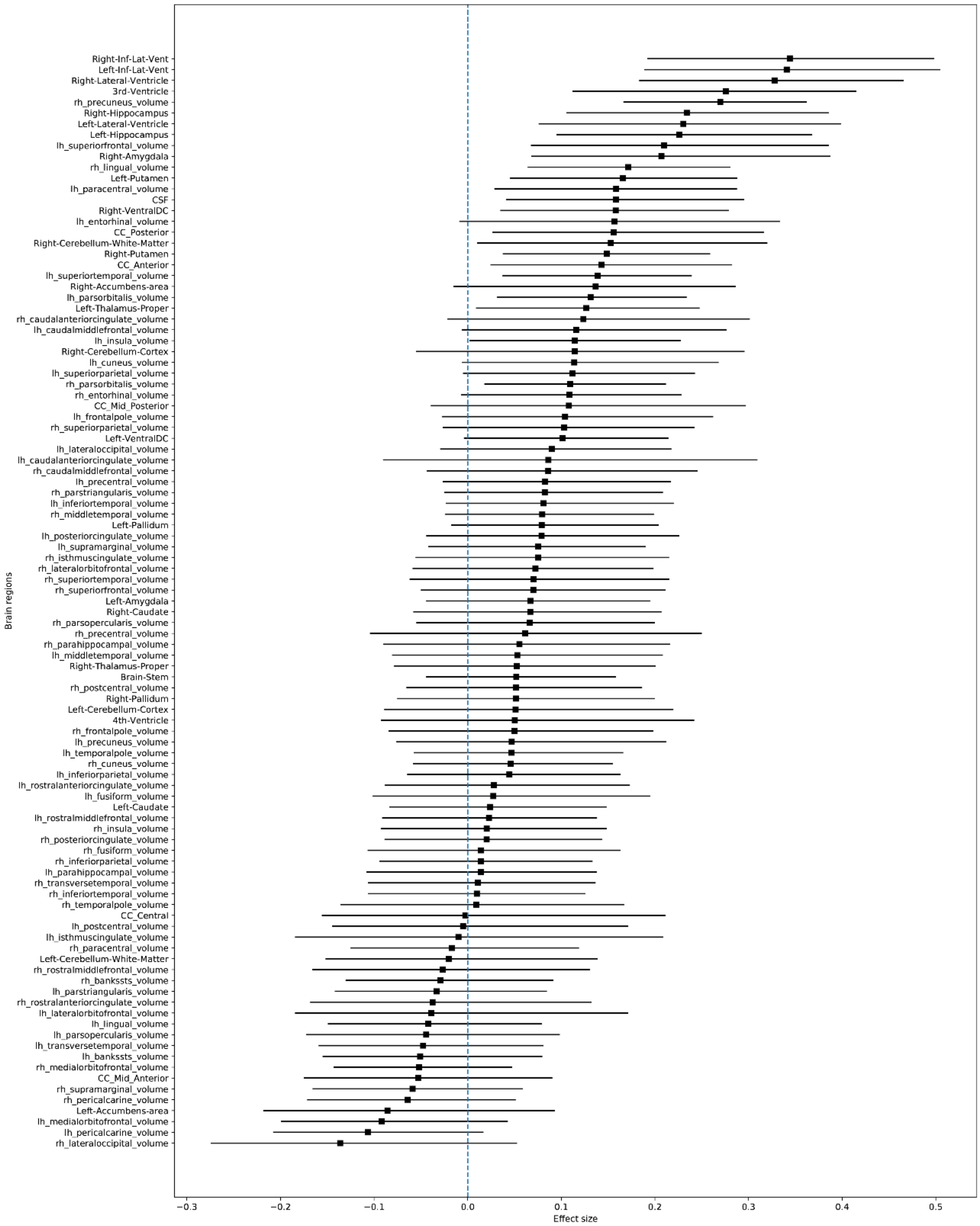


Figure 3 – Regional deviations of the AD group from the ADNI dataset. The marker indicates the mean effect size between the HC and the AD groups. The horizontal bars indicate the 95% confidence interval calculated using the percentile method on the bootstrap analysis.

### 14. Region importance – AIBL dataset – HC vs MCI

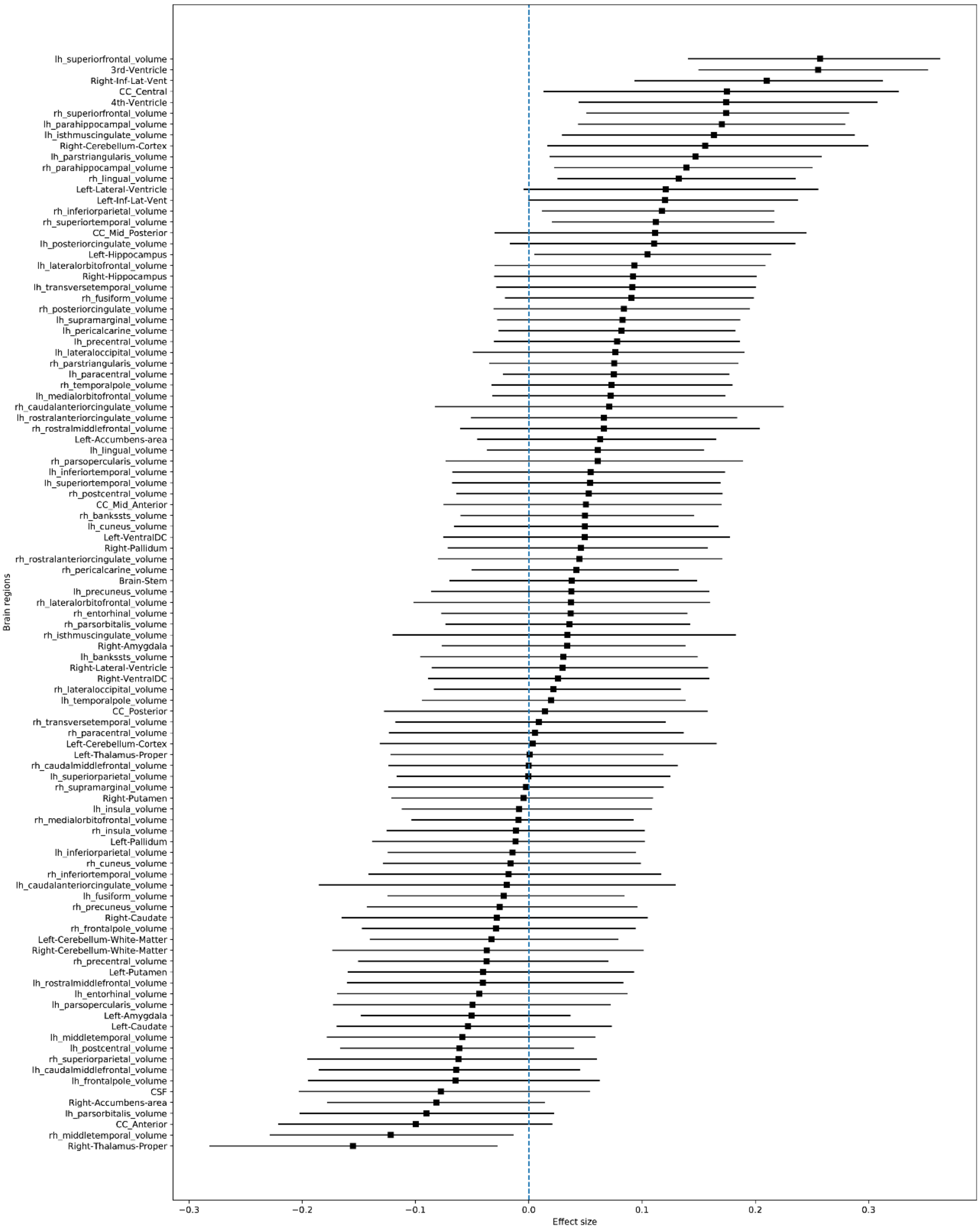


Figure 4 – Regional deviations of the MCI group from the AIBL dataset. The marker indicates the mean effect size between the HC and the MCI groups. The horizontal bars indicate the 95% confidence interval calculated using the percentile method on the bootstrap analysis.

### 15. Region importance – AIBL dataset – HC vs AD

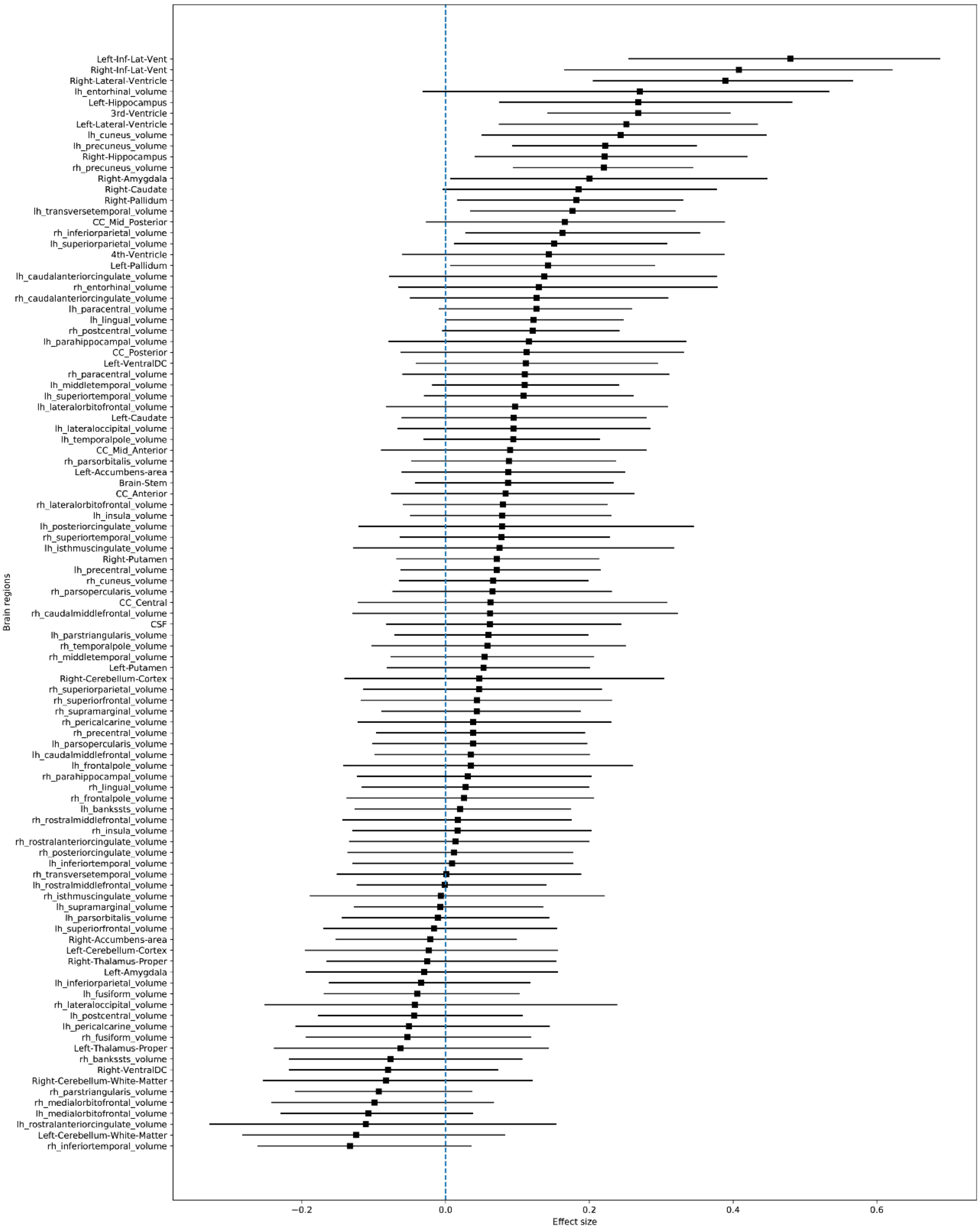


Figure 5 – Regional deviations of the AD group from the AIBL dataset. The marker indicates the mean effect size between the HC and the AD groups. The horizontal bars indicate the 95% confidence interval calculated using the percentile method on the bootstrap analysis.

### 16. Region importance – ARWIBO dataset – HC vs MCI

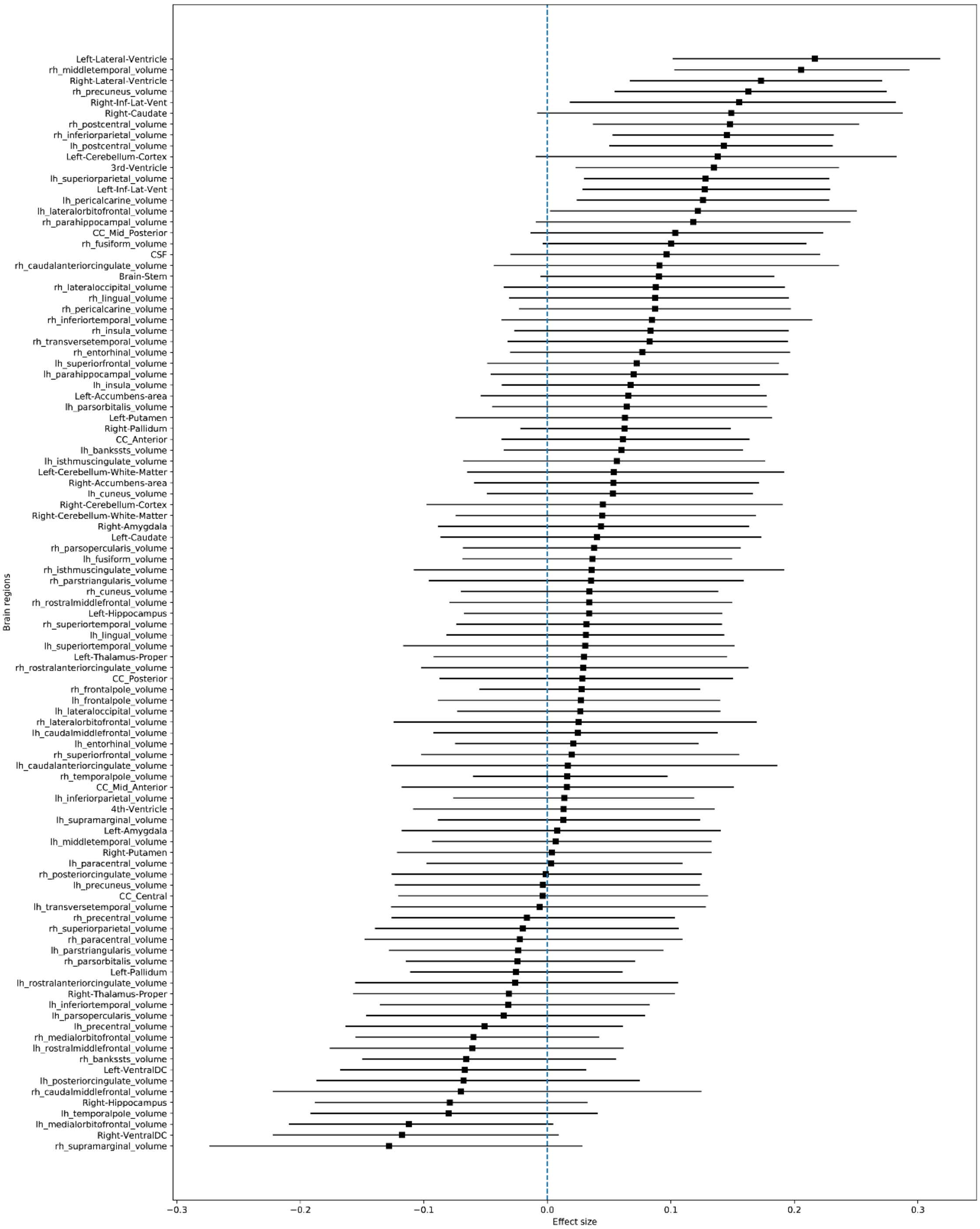


Figure 6 - Regional deviations of the MCI group from the ARWIBO dataset. The marker indicates the mean effect size between the HC and the MCI groups. The horizontal bars indicate the 95% confidence interval calculated using the percentile method on the bootstrap analysis.

### 17. Region importance – ARWIBO dataset – HC vs AD

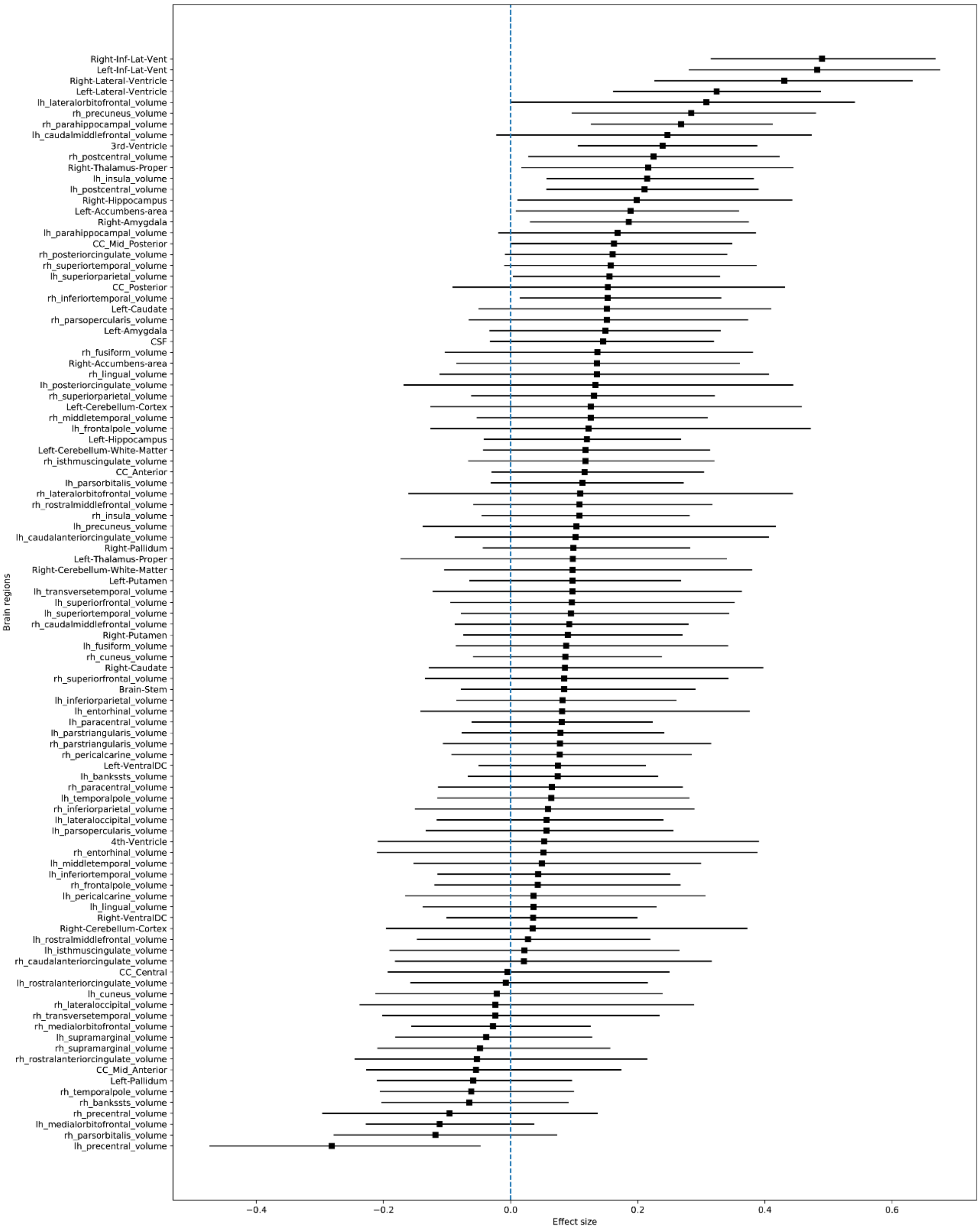


Figure 7 – Regional deviations of the AD group from the ARWIBO dataset. The marker indicates the mean effect size between the HC and the AD groups. The horizontal bars indicate the 95% confidence interval calculated using the percentile method on the bootstrap analysis.

### 18. Region importance – OASIS-1 dataset – HC vs AD

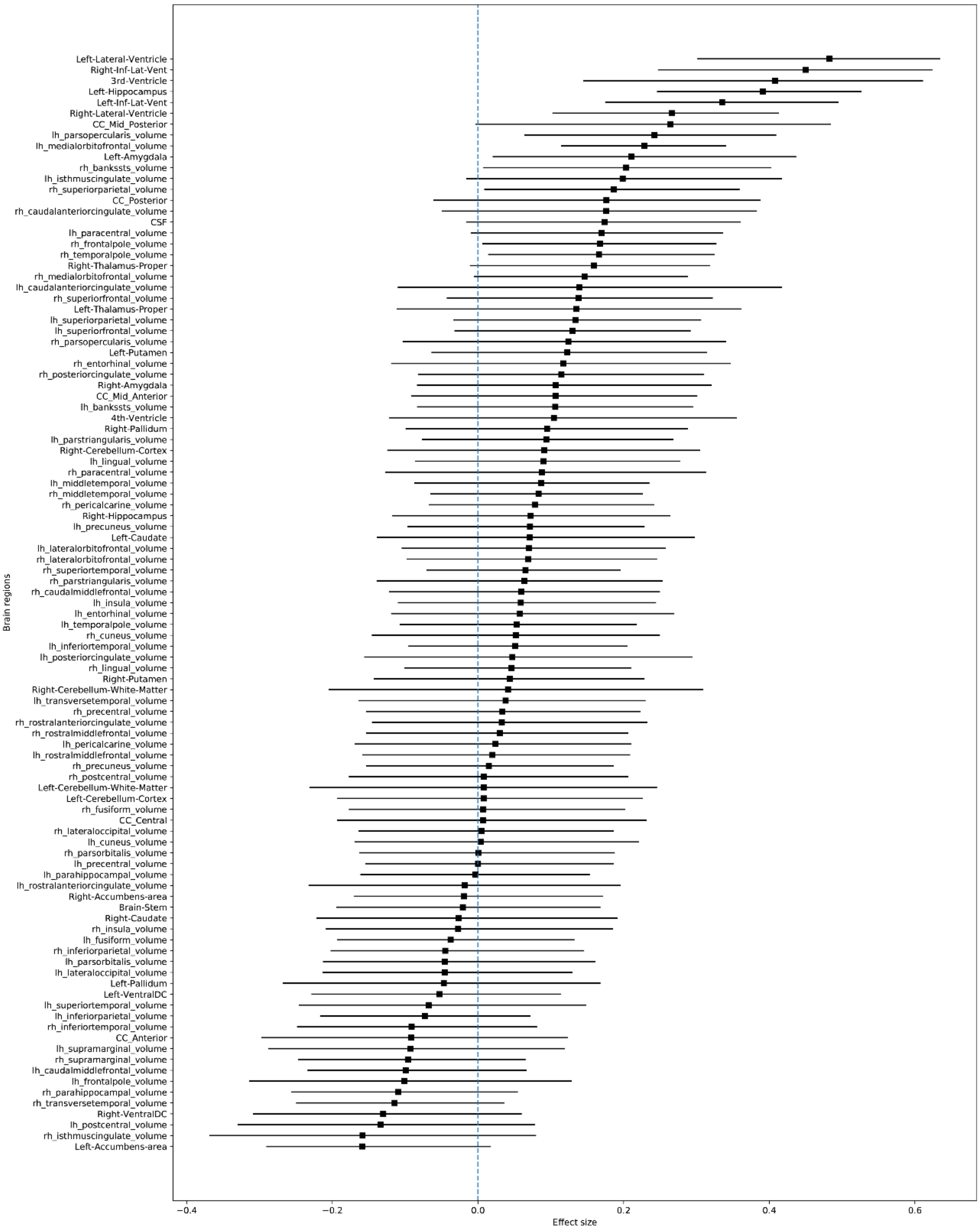


Figure 8 - Regional deviations of the AD group from the OASIS-1 dataset. The marker indicates the mean effect size between the HC and the AD groups. The horizontal bars indicate the 95% confidence interval calculated using the percentile method on the bootstrap analysis.

### 19. Region importance – MIRIAD dataset – HC vs AD

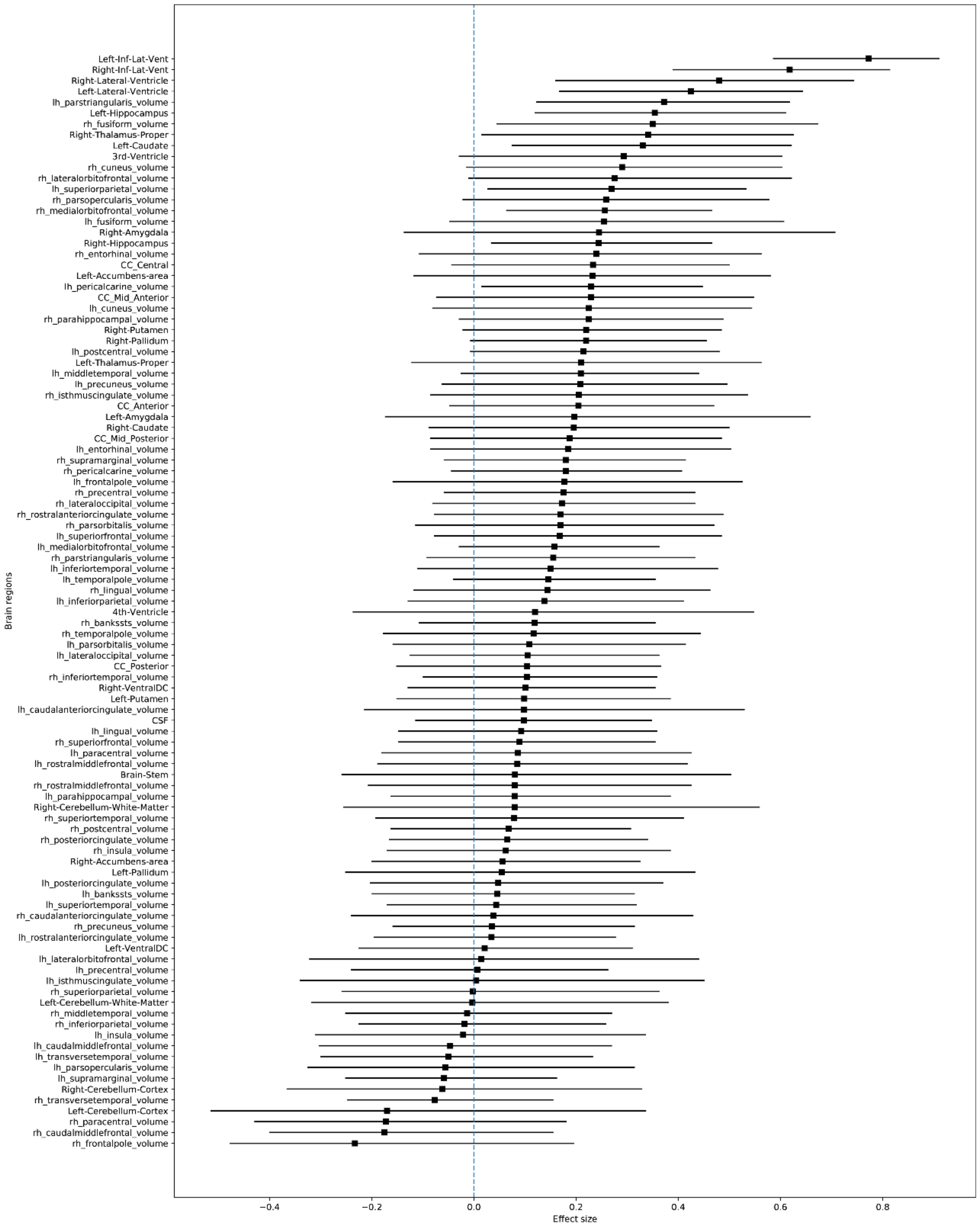


Figure 9 - Regional deviations of the AD group from the MIRIAD dataset. The marker indicates the mean effect size between the HC and the AD groups. The horizontal bars indicate the 95% confidence interval calculated using the percentile method on the bootstrap analysis.
